# VEGFA critically controls neurovascular invasion and chronic low back pain during intervertebral disc degeneration

**DOI:** 10.1101/2025.06.03.657641

**Authors:** Ryan S. Potter, Hong J. Moon, Sade W. Clayton, Remy E. Walk, Addison L. Liefer, Liufang Jing, Evan G. Buettmann, Jennifer A. McKenzie, Alec T. Beeve, Erica L. Scheller, Matthew J. Silva, Amber N. Stratman, Lori A. Setton, Munish C. Gupta, Simon Y. Tang

## Abstract

Despite its enormous burden on patients and society, chronic low back pain (LBP) has no effective therapeutic options. Innervation of the degenerating intervertebral disc (IVD) is suspected to cause discogenic LBP, but the mechanisms that orchestrate the IVD’s neo-innervation and subsequent symptoms of LBP remain unknown. We hypothesize that Vascular Endothelial Growth Factor-A (VEGFA) critically mediates the neurite invasion in the IVD and contributes to chronic LBP. Initiating IVD degeneration through a mechanical injury, we evaluated the progression of neurovascular features into the IVD, as well as ensuring LBP symptoms and locomotive impairments at acute (3-weeks) and chronic (12-weeks) timepoints following the IVD injury. To determine the role of VEGFA, we utilized a mouse model with ubiquitously inducible recombination of the floxed *Vegfa* allele (UBC-Cre^ERT2^; *Vegfa*^fl/fl^). The ablation of VEGFA after an IVD injury attenuated *de novo* neurite and vessel infiltration and impeded the expression of TRPA1, a nociceptive ion channel, in the dorsal root ganglion. The VEGFA-null animals, despite IVD degeneration, exhibited alleviated mechanical allodynia and improved locomotive performance. To determine the effects of IVD-derived VEGFA on endothelial cells and neurons, we co-cultured HMEC-1 endothelial cells and SH-SY5Y neurons with *VEGFA*-silenced human primary IVD cells. The endothelial cells co-cultured with *VEGFA*-silenced IVD cells exhibited reduced vessel growth and shifted their transcriptome and secretome from angiogenic to lymphangiogenic. The neurons co-cultured *VEGFA*-silenced IVD cells showed slowed growth and attenuated transcriptional programs for growth and elongation. These results show that VEGFA directs the growth of intradiscal vessels and neurites that cause low back pain and impaired function, and the inhibition of IVD-derived VEGFA during degeneration may be sufficient to prevent chronic pain behavior and motor impairment associated with discogenic low back pain.

**One Sentence Summary:** VEGFA is a key mediator of neurovascular infiltration in the degenerating intervertebral disc and an essential driver of chronic low back pain, whose ablation prevents pain-related behaviors.

## INTRODUCTION

Low back pain (LBP) is a prevalent, debilitating, costly, and multifactorial condition worldwide, imposing substantial global and economic burdens^1–3^. LBP affects as much as 80% of the population, and the elderly experience greater rates of LBP that exacerbate their functional decline, frailty, and loss of independence^4^. LBP is the leading cause of years lived with disability and contributes an estimated $50 to $100 billion USD annually in healthcare costs^4–6^. Chronic LBP is commonly associated with intervertebral disc (IVD) degeneration, which occurs with aging, genetic predisposition, and injury^7–10^. Degeneration leads to structural and biochemical changes in the IVD, including the loss of hydration, inflammation, and matrix breakdown^11–14^. Despite our understanding of the features of IVD degeneration, the specific degenerative features that are mechanistically responsible for the eventual low back pain remain unknown^15^. While the healthy IVD is relatively aneural, the painful human intervertebral discs are innervated, suggesting a role for aberrant nerve growth in discogenic pain^16^. VEGFA (Vascular Endothelial Growth Factor A) is a potent signaling protein that promotes the proliferation, migration, and survival of endothelial and neural cells^17–19^. While VEGFA has known roles in both vascular and neural support, its function following an IVD injury, and how it initiates or sustains discogenic pain remains unresolved. We hypothesized that VEGFA is crucial for the neurite growth and vascularization in the injured IVD to mediate acute and chronic discogenic pain.

To investigate the role of VEGFA in IVD degeneration and low back pain, we utilized a mouse model with an inducible global ablation of VEGFA. To model discogenic low back pain, we created an injury in the lumbar IVD to initiate degeneration^20–22^. Following the IVD injury, we administered tamoxifen to induce the ablation of the *Vegfa* gene. We subsequently measured pain-related symptoms such as mechanical and thermal sensitivity as well as locomotor function^20^. Because VEGFA is also a critical regulator of vascular growth, we assessed neo-vascularization into the injured IVD along with neural infiltration to better understand the relationship between vascular and neural response in disc degeneration. To evaluate whether there was peripheral nervous system adaptation, we evaluated pain-related ion channels expression in the lumbar dorsal root ganglia (DRGs). We performed co-cultures of human cells to determine the role of IVD-produced VEGFA on neuron and endothelial cell growth.

Our findings show that VEGFA plays a central role in promoting neurovascular ingrowth into the degenerating intervertebral disc. The ablation of VEGFA reduced neurite and blood vessel infiltration into the degenerated IVD and alleviated the mechanical allodynia and locomotor deficits in mice. VEGFA ablation also attenuated pain-related ion-channel expression in the DRGs of the injured animals. Diminished endothelial and neuronal growth *in vitro* with VEGFA-silenced IVD cells highlights the key role of IVD cell derived VEGFA in promoting vascular and neurite infiltration. These results demonstrate that targeted inhibition of VEGFA can suppress intradiscal vessels and neurites that cause peripheral system sensitization, low back pain symptoms, and impaired function.

## RESULTS

### Removal of VEGFA Blunts Intradiscal Invasion of Neurovascular Structures

In WT (Cre−;*Vegfa*^fl/fl^ with tamoxifene administration) animals, the injured IVD was infiltrated by PGP9.5+ neurons and by Endomucin+ vessels. VEGFA ablation significantly blunted these neuronal and vascular features. Immunofluorescence images from 50µm thick sections (Figure 1A-B) showed fewer neural and vascular structures in VEGFA-null mice compared to WT particularly in the region around the outer annulus fibrosus (AF) near the injury site. In a validation study, at least 94% of the *de novo* PGP9.5 positive structures within the IVD were positive for Na_v_1.8 (Figure S3), a marker for sensory neurons^60^. Quantitative measurements revealed that VEGFA removal diminished neurite length (p=0.01; Mean difference = -57.34, 95%CI:-103 -12.1[μm]), vessel length (p=0.004; Mean difference = -177, 95%CI:-295 -59.4[μm]), neurite penetration depth (p=0.002; Mean difference = - 15.25, 95%CI: -24.9 -5.72[μm]), and vessel penetration depth (p=0.047; Mean difference = -18.80, 95%CI:-37.3 -0.284[μm]) (Figure 1C-F). Post hoc analysis reveals differences between the WT and VEGFA-null IVDs in neurite depth at 3 weeks (p=0.001) and in vessel length at 12 weeks (p=0.01). These findings show that VEGFA is crucial for instigating the neurite growth and sustaining vessel propagation in IVDs following injury. No sex-differences were observed (Table S1).

**Fig. 1:**
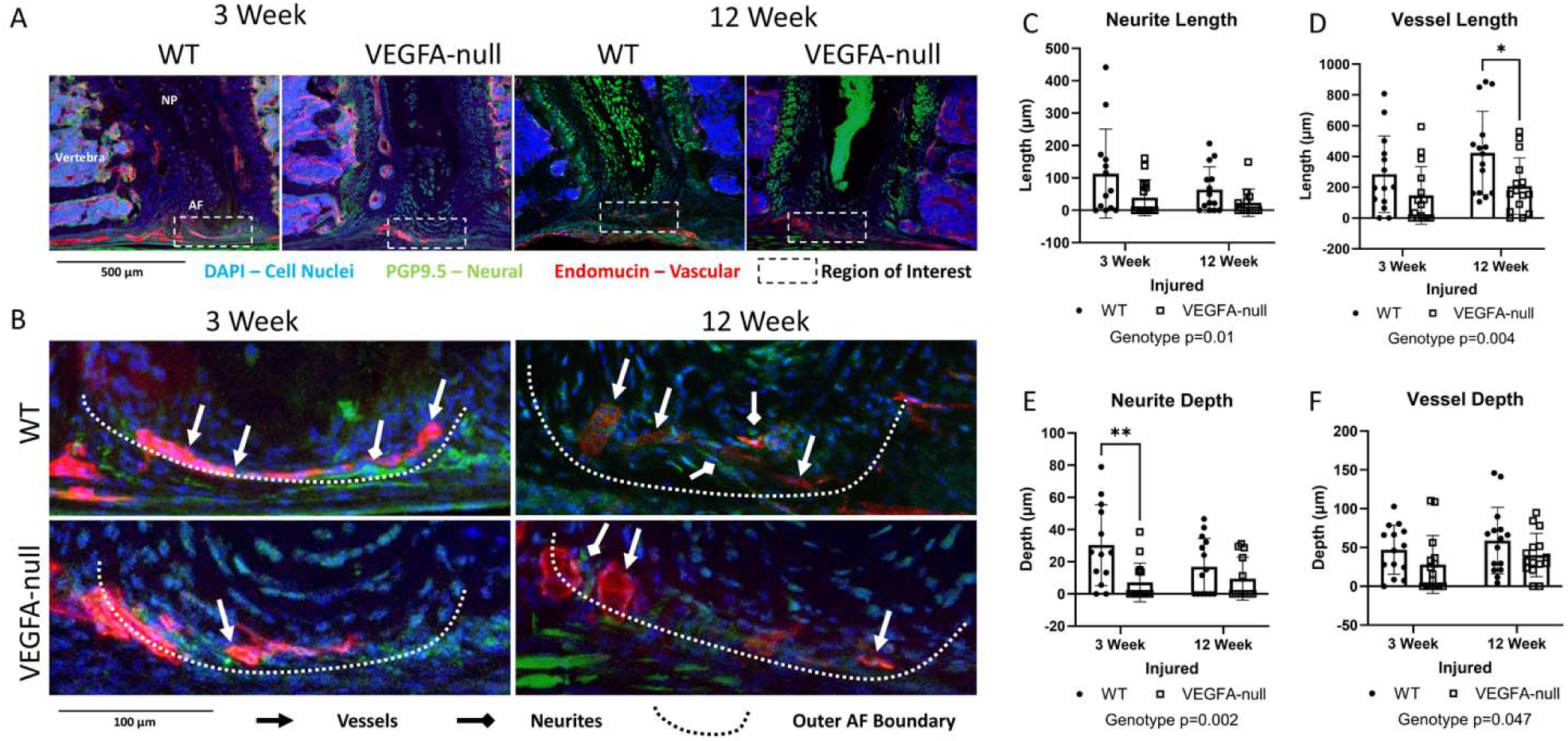
Removal of VEGFA Blunts the Invasion of Neurovascular Structures on the Intervertebral disc dorsal surface Following lumbar IVD Injury. A-B) IHC with the nuclei (DAPI) in blue, neuronal features (PGP9.5) in green and vascular features (Endomucin) in red. C-F) VEGFA-null animals had diminished neurite lengths (p=0.01; Mean difference = -57.34, 95%CI:-103 -12.1[μm]) and vessel lengths (p=0.004; Mean difference = -177, 95%CI:-295 -59.4[μm]). Likewise, the neurite penetration depth (p=0.002; Mean difference = -15.25, 95%CI: -24.9 - 5.72[μm]) and vessel penetration depth depth (p=0.047; Mean difference = -18.80, 95%CI:-37.3 - 0.284[μm]) are blunted by VEGFA removal. Post hoc analysis reveals differences between the WT and VEGFA-null IVDs in neurite depth at 3 weeks (p=0.001) and in vessel length at 12 weeks (p=0.01). Mean ± standard deviation are shown in the graphs.

### VEGFA Is Essential for the Spatial Colocalization of Intradiscal Nerves and Vessels

The spatial proximity of intradiscal nerves and vessels are disrupted by VEGFA ablation. Immunofluorescence images reveal the distribution of neural (PGP9.5, green) and vascular (Endomucin, red) structures highlighting the regions of interest around the outer annulus fibrosus (AF) and overlapping lengths relative to neurites (Figure 2A-B). The colocalization analysis shows that the WT injured IVDs have neurites with a large amount of vessels in their proximity. The removal of VEGFA in the IVDs 3 weeks following injury protected more IVDs from neurites penetration (Figure 2C;p =0.02; OR=4.275, 95%CI:1.44 12.6). The loss of VEGFA also protected more IVDs from the penetration of endomucin+ vessels (Figure 2D; p=0.03; OR=10.67, 95%CI:1.69 122). The spatial overlap analysis between neurites and vessels shows that VEGFA ablation dramatically reduces the amount of neurites near vessels (Figure 2E; p=0.007; Mean difference = -48.8, 95%CI: -84.0— - 13.69[μm]). After adjusting for amount of neurites in each animal, this observation remains robustly independent of the basal level of neurites (Figure 2F p=0.008; Mean difference=-36.1, 95%CI: - 0.943—-71.2[μm]). The lack of difference over time in the depth and the length ratios between the neurites and vessels indicates that once initiated, VEGFA does not appear to affect the propagation of these features (Figure 2G-H). These findings highlight the critical role of VEGFA in the *de novo* initiation of neural and vascular features in the IVD.

**Fig. 2:**
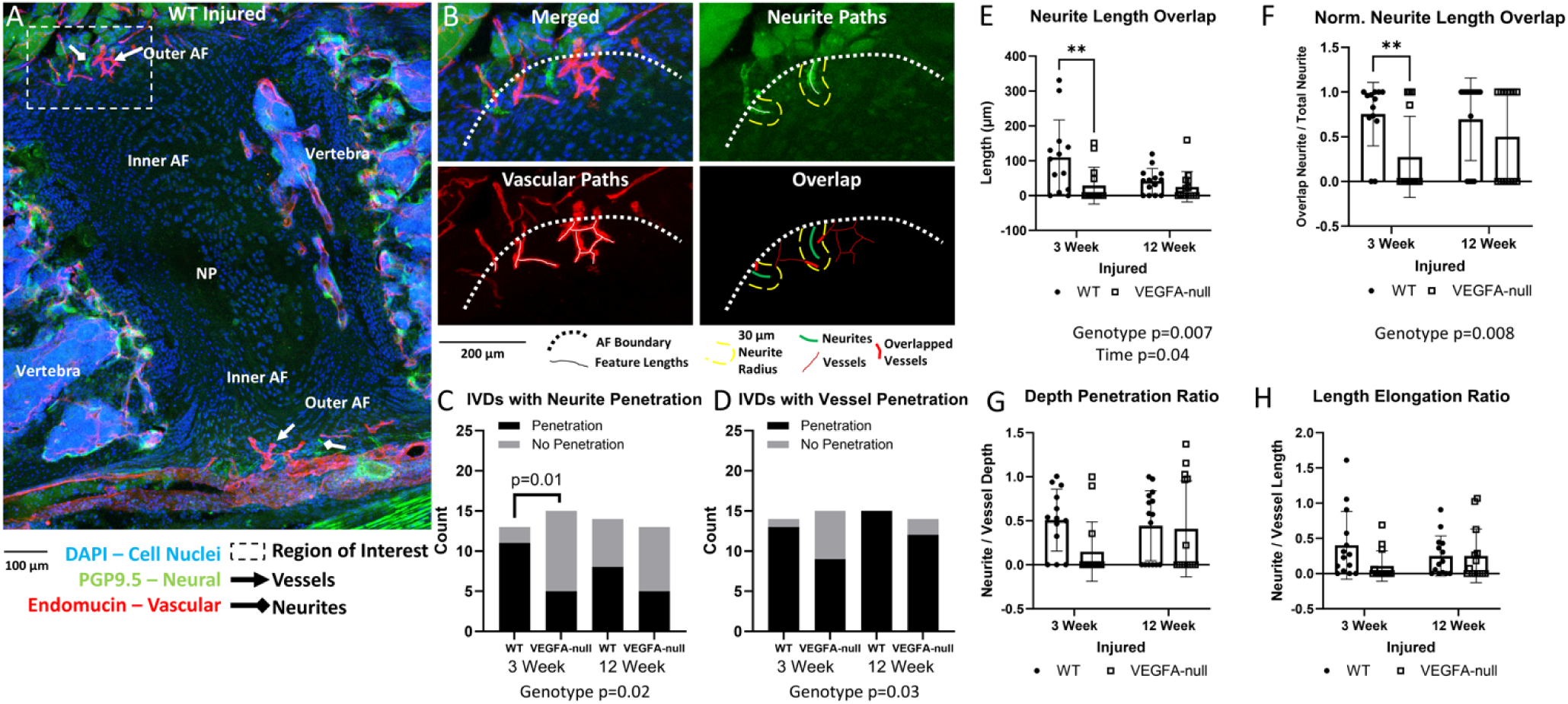
VEGFA is Essential for the Colocalization of Intradiscal Nerves and Vessels and for *de novo* neurovascular invasion. A) An injured WT IVD and the ROI for the analysis neurovascular features. B) Graphical representation of how spatial colocalization analyses quantified the number of neurites that are with a 30-µm proximity of vessels. C-D) Neurite (p =0.02; OR=4.275, 95%CI:1.44 12.6) and vessel (p =0.03; OR=10.67, 95%CI:1.69 122) infiltration proportions were both significantly higher in WT animals than in the VEGFA-null groups. E) VEGFA ablation suppressed vessel-adjacent neurites (p=0.007; Mean difference = -48.8, 95%CI: -84.0—-13.69[μm]). F) The loss of VEGFA prevented neurite-vessel spatial coupling even when accounting for varying intradiscal neurite lengths (p=0.008; Mean difference=-36.1,95%CI: -0.943—-71.2[μm]), confirming that this phenomenon is conserved even when accounting for heterogenous vessel sprouting across animals. G-H) The lack of an effect of time in the depth of penetration and length elongation ratios suggest that once the neurites and vessels have initiated, the loss of VEGFA does not appear to affect their propagation. Statistical model C-D: Fisher’s exact test. Statistical model E-H: 2 factor ANOVA with Fisher LSD post-hoc comparisons. E-H Data are presented as mean ± standard deviation.

### Depletion of VEGFA Suppresses the Expression of Nociceptive Ion Channels in the DRG

Quantitative measurements of corrected total cell fluorescence (CTCF), averaged from the left and the right sides, of the L2-L3 lumbar dorsal root ganglia (DRGs; Figure 3B-D) revealed a significant reduction in transient receptor potential ankyrin 1 (TRPA1) fluorescence (Figure 3D; p=0.037;Mean difference=-16040, 95%CI: -30800—-1280) due to VEGFA removal following injury. No significant changes were measured in tyrosine kinase receptor A (TrKA) or transient receptor potential vanilloid 1 (TRPV1) CTCF levels due to the loss of VEGFA ablation.

**Fig. 3:**
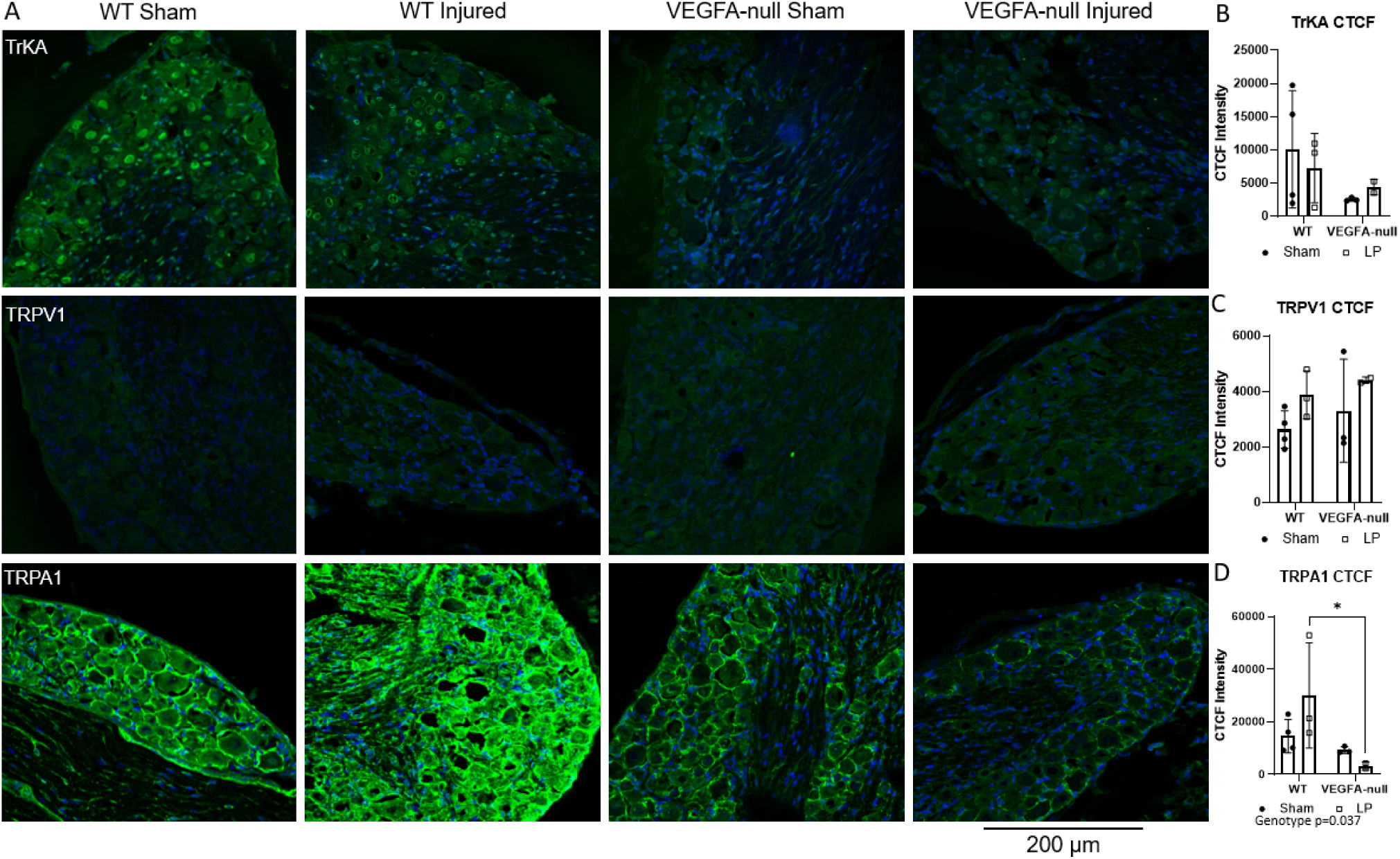
VEGFA Inhibition Reduces the Expression of Pain-related Ion Channels in the DRG. A) TrKA (top row), TRPV1 (middle row), and TRPA1 (bottom row) in green and DAPI in blue were quantified. B-C) No effects due to VEGFA were observed in TrKA or TRVP1 expression measured by Corrected total cell fluorescence (CTCF). D) VEGFA ablation suppressed TRPA1 expression (p=0.037;Mean difference=-16040,95%CI:-30800 — -1280). TRPA1 expression is canonically associated with nociceptive sensitization.

### Ablation of VEGFA Alleviates Mechanical Sensitivity and Locomotive Function

#### Loss of VEGFA Attenuates Mechanical, but Not Hot or Cold, Sensitivity

WT animals exhibited mechanical sensitivity measured by the Von Frey filament test at both 3 and 12 weeks after the IVD injury compared to the pre-surgery baseline. Ablation of VEGFA prevented this impairment (Figure 4A; p=0.01, Mean difference=0.490, 95%CI: 0.110—0.860). The loss of VEGFA reversed the mechanical allodynia at 12 weeks. The electronic Von Frey filament test also revealed that there is a greater mechanical sensitivity on the side the surgical approach (e.g. left). VEGFA removal specifically alleviated the mechanical sensitivity of the injured side, without modifying the contralateral side (Figure S1G). No notable changes were observed in heat or cold sensitivity at any post-operative time point (Figure 4B-C; Table S1).

**Fig. 4:** VEGFA Ablation Improves Mechanical Sensitivity and Performance. A) The loss of VEGFA mitigated the mechanical sensitivity of the animals with injured IVDs (p=0.01;Mean difference=0.490, 95%CI: 0.110—0.860). In particular, the VEGFA-deficient animals recovered from the initial mechanical sensitivity at 9- and 12- weeks while the WT animals do not recover (p<0.01). B-C) No differences in hot or cold sensitivity were observed. D) There were significant differences in longitudinal measures of performance that included rotarod (genotype, p=0.004; time, p=0.003), indicative of a time-dependent VEGFA effect on the progression of performance. E-F) There was no differences in any measures of open field with genotype or time point. G-I) There was no differences in the inverted screen endurance or hind paw grip strength with genotype and at any time point. Statistical model A-F, I: Linear mixed effects with genotype and time as factors with post-hoc LSM with Sidak correction. G-H Mantel-Cox test. A-F, I Data are presented as mean ± standard deviation.

#### VEGFA Ablation Improves Rotarod Performance, but Does Not Affect Passive Function

We longitudinally evaluated locomotive performance through an active challenge of physical function, measured through the rotarod assay, every three weeks up to 12 weeks after injury. The longitudinal rotarod test reveals that VEGFA-null mice exhibited longer latencies to fall compared to WT (Figure 4D; p=0.004, Mean difference=15.13, 95%CI:5.35—24.9), in contrast to the progressive decline of motor coordination over time in the WT animals (p=0.003). Remarkably, at 9- and 12- weeks, the initial decline of rotarod performance in VEGFA null animals was recovered. In the Open Field test, total distance traveled (Figure 4E) and rearing counts (Figure 4F) were not different between genotypes, sex, tamoxifen, or due to surgery (Table S1), suggesting comparable levels of exploratory behavior and general activity between all groups.

#### VEGFA Removal Does Not Affect Measures of Strength

To determine the impacts on musculoskeletal function following VEGFA ablation, strength assessments were conducted, including inverted screen endurance and hind paw grip strength. The inverted screen endurance test, presented using Kaplan-Meier curves, showed that VEGFA-null mice have similar probabilities of survival (e.g. hanging on the wired mesh) compared to WT at both 3 weeks (Figure 4G) and 12 weeks (Figure 4H). Measures of hind paw grip strength (Figure 4I) were also found to be similar across genotypes, indicating no compromise in muscle function from VEGFA ablation.

Overall, the behavioral and locomotive function analyses show that VEGFA-null mice have improved recovery of mechanical sensitivity. Further, the VEGFA-null animals averted the transition to chronic sensitivity, while also avoiding the performance deficits observed in WT at the chronic time points.

### Ablation of VEGFA does not affect structure, function, or degeneration of the IVD

#### Loss of VEGFA Does Not Affect the Histopathologic Degeneration of the IVD

CEµCT analysis of the IVDs 3 weeks after injury reveals significant deterioration of the IVD structure. By using a contrast enhancing agent, Ioversol, the injury was observed on the left ventral side of the outer AF as progressively higher attenuating regions that mark the fibrous scar tissues (Figure 5A). The injured IVDs also exhibit a loss of NP attenuation, measured by NI/DI, demonstrating the loss of the water retaining ability of the NP (Figure 5B; p=0.03,Mean difference=-0.0102,95%CI:-0.0190—-0.00138). The ablation of VEGFA does not affect injury-mediated degeneration. A narrowing of the intradiscal space measured by disc height ratio (DHR), a hallmark of IVD degeneration, is observed in all of the injured discs (Figure 5C; p=0.01,Mean difference=-0.0246,95%CI:-0.0431—-0.00605) with no differences among genotypes or with post-operative time. There was not a change in NP volume either due to injury or VEGFA ablation (Figure 5D). Injury increases the degenerative features including decreased NP hydration and a narrowing DHR; VEGFA ablation did not appear to alleviate these tissue-level changes in the IVD.

**Fig. 5:**
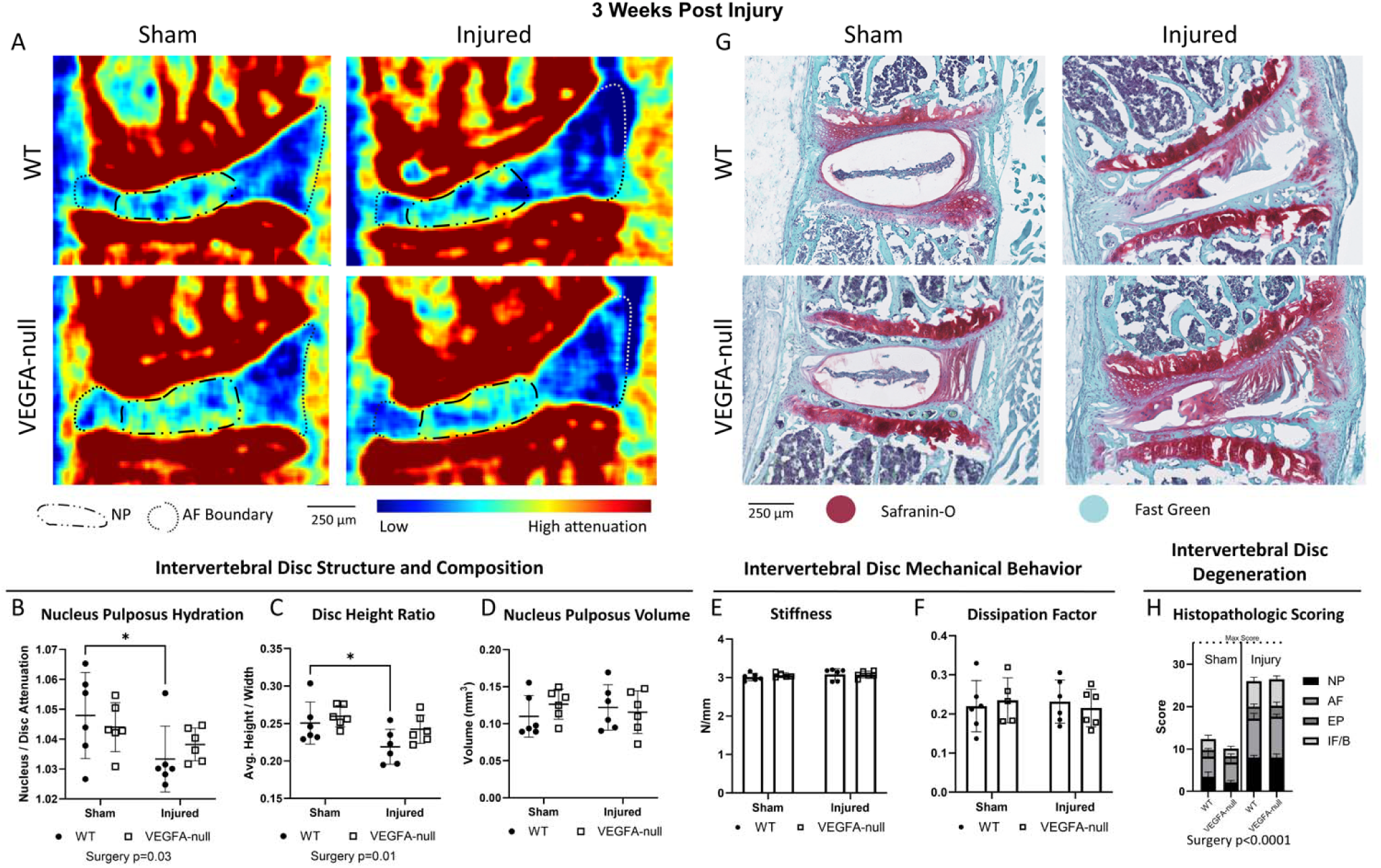
Loss of VEGFA Does Not Alter Structural, Mechanical, or Histopathologic Degeneration of the Injured IVD. A) IVD structure analyses via CEµCT show that B) injury decreased NP hydration (p=0.03, Mean difference=-0.0102, 95%CI:-0.0190—-0.00138), indicated by the decreased attenuation of the NP compared to sham. C) Injury decreased the IVD’s disc height ratio, DHR, (p=0.01, Mean difference=-0.0246, 95%CI:-0.0431—-0.00605) compared to sham. D) No effect of injury was observed on NP volume. Changes to mechanical behavior were not observed in E) stiffness and F) dissipation factor, indicating that tissue level mechanics were not changed by injury or VEGFA removal at 3-weeks post injury. G) Histological images stained with Safranin-O and Fast Green show IVD degeneration. H) Histopathologic scoring shows a strong effect of injury (p<0.0001;Mean difference=15.2, 95%CI:13.5—16.5) with no statistically distinguishable differences due to VEGFA loss in nucleus pulposus, annulus fibrosus, end plate, or the interface/boundary demonstrating the pathophysiological response from needle puncture. Data are presented as mean ± standard deviation.

#### Injury to the IVD Does Not Impair Its Mechanical Function

Dynamic compression of the IVDs 3 weeks post injury shows no distinct mechanical responses to injury or impact from VEGFA ablation. The stiffness measurements show no significant differences between WT and VEGFA-null groups, both in sham and injured conditions (Figure 5E). Likewise, the dissipation factor indicates no appreciable changes from injury or VEGFA loss (Figure 5F). These findings show that the degeneration that occurs 3 weeks after injury in this model does not yet impair the IVD’s mechanical function.

#### VEGFA Loss Does Not Rescue the Degenerative Changes Caused by Injury

Histopathological scoring (Figure 5G-H) revealed the damaging effects of the needle stab injury on the compartments in the disc, including decreased NP cell density and changes in cellular morphology, disruption of the AF concentric lamellar structures with increased cellularity and presence of clefts, along with thinning and disruptions in the endplate cartilage, and structural disorganization in the boundary regions. Scores for injured groups averaged near 26 out of a possible 36 for maximal degeneration, while sham controls averaged 11 (Figure 5G,H; p<0.0001; Mean difference=15.2, 95%CI:13.5—16.5). Effects of VEGFA removal were not detected. IVD injury dramatically changed all compartments of the disc with the largest effects on the AF followed by NP while sham scores indicate mild degeneration from the surgical exposure of the disc and likely disruption of surrounding soft tissues.

### Silencing *VEGFA* RNA in Human IVD Cells Attenuates the Growth of Co-Cultured Neuronal Cells (NC) and Endothelial Cells (ECs)

Transfecting human IVD cells with VEGFA-specific small interfering RNA (siRNA) suppressed their IL-1β-stimulated VEGFA production by 57.4% (Figure S6). After removing the IVD cells from the IL-1β laden media, co-culturing these VEGFA-siRNA transfected IVD cells with SH-SY5Y neuroblastoma-derived neuronal cells (aka NCs) maintained the NCs’ morphology but blunted the axonal growth (Figure 6A). The total neurite length reveals declined neurite growth in NCs co-cultured VEGFA-siRNA transfected IVD cells (Figure 6B-D). RNA sequencing revealed that NCs co-cultured with VEGFA-knockdown IVD cells differentially downregulated 39 genes and upregulated 107 genes (aka DEGs) compared to those co-cultured with VEGFA-intact IVD cells (Figure 6E). Gene ontology analysis determined the most enriched biological process from the DEGs were involved GPCR signaling pathways, known for neurovascular development (*EDNRA, CHGA, RAMP3*), regulation of growth (*DCC, GREM1, STC2, MYOZ1, MT1F, BST2, IGFBP3, CXCR4, MT1X, IGFBP7, BCL2*), and the negative regulation of cell population proliferation (*GREM1, PTPN14, DHRS2, SSTR2, SCG2, SERPINF1, IGFBP3, OVOL1, ADM, IGFBP7, BCL2, S100A11*). Notably, several DEGs were specific to cell growth inhibition (*SSTR2, S100A11),* immune cells and inflammation (*CXCR4, FGFB2, IL32)*, and neuronal cell functions (*DCC, SDK2*, *ASTN2*).

**Fig. 6:** Co-culturing SH-SY5Y neuronal cells with VEGFA-silenced human primary IVD cells suppressed *in vitro* neurite growth and transcriptional processes related to growth and elongation. A-D) The neurite extension assay after co-coculturing neuronal cells with IL-1β stimulated VEGFA siRNA transfected human IVD cells shows reduced total neurite lengths. E) Volcano plot of RNA-seq analysis showing 146 differentially expressed genes and F) key biological processes regulated by VEGFA siRNA. NCs co-cultured with IL1β naïve disc cells showed undifferentiated cells possessing few short projections which cluster together, while NCs co-cultured with IL1β+SCRAM siRNA and NCs co-cultured with IL1β+VEGFA siRNA showed differentiated cells having many extensive projections.

We also co-cultured VEGFA-siRNA transfected IVD cells, stimulated a priori with IL-1β, with the HMEC-1 endothelial cells (aka ECs). The ECs co-cultured with VEGFA-silenced human IVD cells formed fewer closed tubes, a measure of vessel formation, after 6 hours of culturing (Figure 7A-B; p=0.04). RNAseq uncovered 175 downregulated and 0 upregulated DEGs when ECs are co-cultured with IVD cells transfected with VEGFA siRNA compared with scram (Figure 7C). Gene ontology analysis determined the most enriched biological processes were involved in vessel and vasculature related processes. These included lymphatic vessel development (*NR2F2, TBX1, SOX18, EFNB2, HEG1*), blood vessel morphogenesis (*TNFAIP2, EFNB2, HEG1, PRKACA, NRARP, TBX1, SOX4, NR2F2, AKT1, QKI, SMAD7, SOX18, VEGFB*), and regulation of neuron projection development (*EFNB2, CDK5R1, RYK, RAPGEF1, CARM1, AKT1, PLZNA1, NCS1, DVL1, SLK, STK24, BRSK2, ADNP*). Interestingly, the canonical WNT signaling pathway was also an enriched biological pathway. WNT signaling plays an important role in endothelial cell proliferation, survival, differentiation and response to mechanical forces due to shear stress from blood flow.^23,24^

**Fig. 7:**
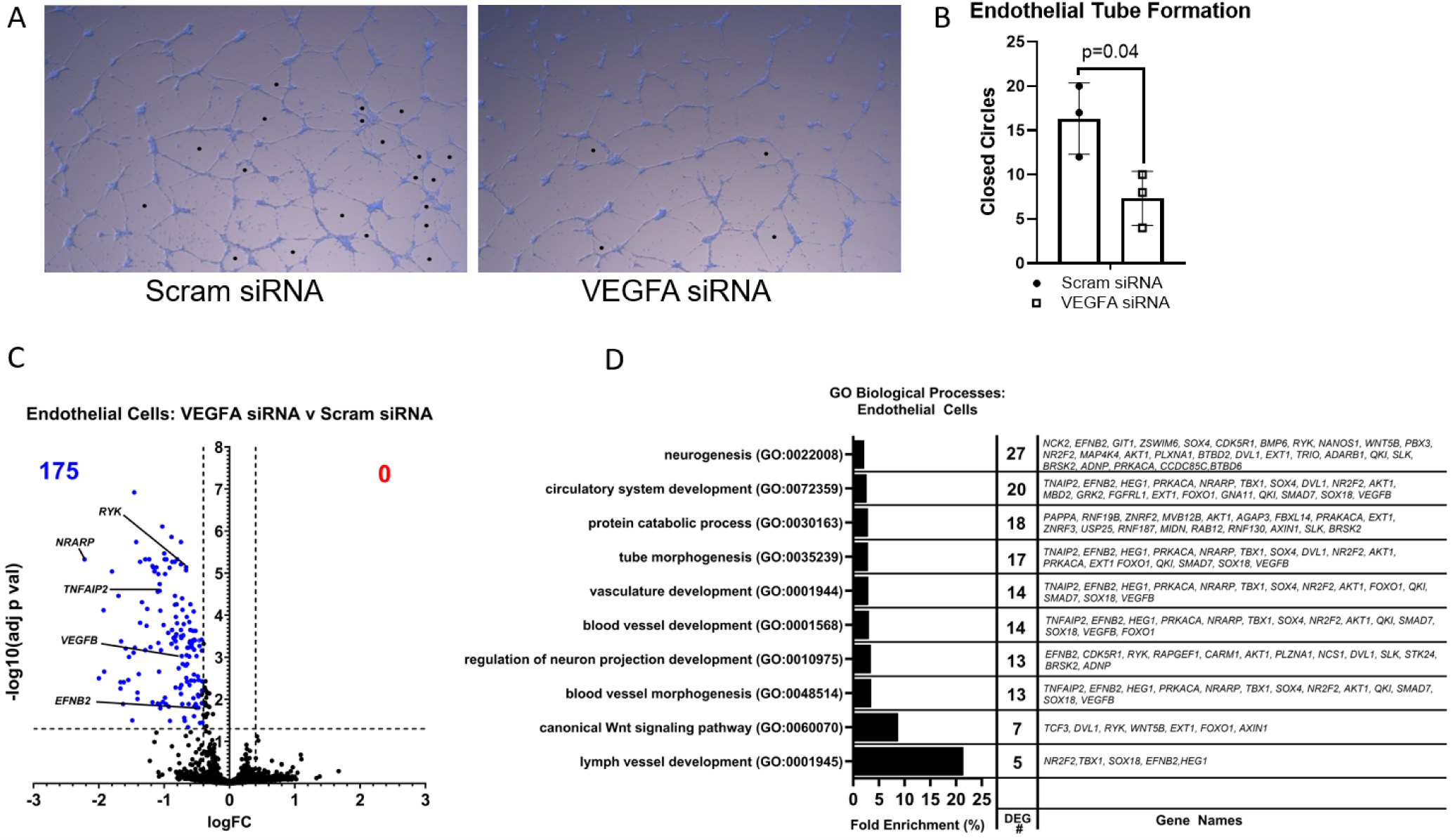
Co-culturing with VEGFA-silenced human primary IVD cells suppressed *in vitro* endothelial cell growth and shifts the endothelial cell transcriptional program from angiogenic to lymphangiogenic. A) Matrigel tube forming assay images of endothelial cells (ECs) cocultured with human IVD cells treated with siRNA showing initiation of vessel formation after 6 hours. B) The silencing of VEGFA siRNA in IVD cells reduced tube formation in EC co-cultures, compared to IVD cells with Scrambled siRNA (p=0.04). C) The ECs co-cultured with VEGFA-silenced IVD cells downregulated 176 genes. Key angiogenic genes were suppressed including *HEG1*, *NARAP*, and *NCS1*. There is also a suppression of endothelial cell-derived neurogenic genes including *BRSK2* and *CRLF1*. D) The RNAseq analyses of the HMEC-1 also show an ontological shift to lymphangiogenesis, matrix remodeling, and suppression of neurovascular regulating pathways following co-culture with VEGFA-silenced IVD cells.

#### The endothelial cells (ECs) co-cultured with the VEGFA-silenced IL-1β-stimulated IVD cells produce less angiogenic and inflammatory factors and more non-VEGFA proangiogenic compensatory factors

Following co-culture with VEGFA-silenced, IL-1β-stimulated IVD cells, the ECs exhibited a distinct cytokine profile that indicate a shift in cellular function. Notably, there was a reduction in secretion of VEGFA (Figure 8A) and CCL3 (Figure 8K) protein in the ECs co-cultured with VEGFA-silenced IVD cells, indicating decreased angiogenic and inflammatory signaling. The ECs exposed to VEGFA-silenced IVD cells also produced more VEGFC protein, a canonical driver of lymphangiogenesis and lymphatic endothelial cell proliferation^25,26^. Simultaneously, ECs increased protein production of PDGF-AA (Figure 8J), a mitogenic factor that can compensate for VEGFA loss by promoting vessel stabilization and pericyte recruitment^27^. Increases in MMP1, MMP3, MMP10, and MMP13 protein levels (Figure 8C-F) reflect heightened extracellular matrix remodeling, a process essential for vessel pruning and structural reorganization typically associated with either vessel regression or tissue remodeling rather than active angiogenesis^28,29^. The elevation of granulocyte colony-stimulating factor (G-CSF) and granulocyte-macrophage colony-stimulating factor (GM-CSF) supports a role for immune cell recruitment and activation, especially of myeloid lineages that modulate inflammation and tissue repair (Figure 8G-H)^30,31^. Similarly, upregulation of CXCL1 and CCL5 (RANTES) reinforces the potential for leukocyte chemotaxis and immune surveillance (Figure 8I,L)^32,33^.

**Fig. 8:**
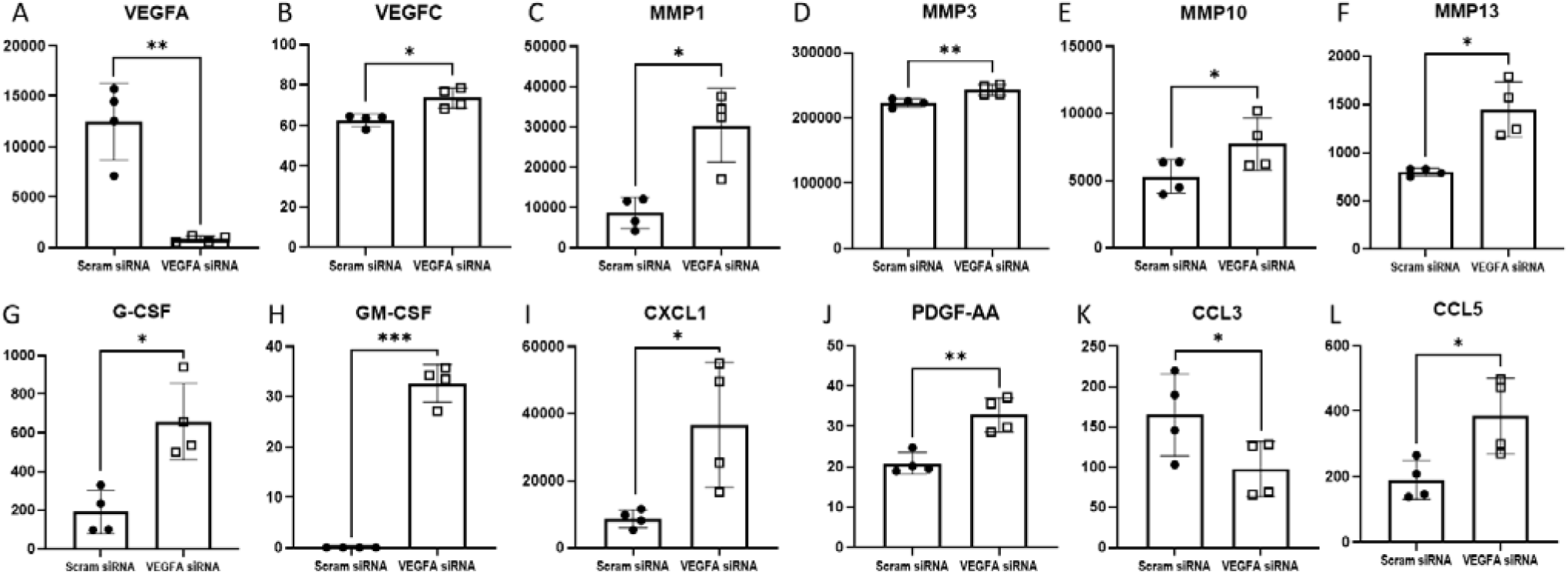
The secreted factors from HMEC-1 (aka ECs) co-cultured with IVD cells showed compensatory factors for angiogenic homeostasis and shifts to lymphanogenesis. Despite having a functional *VEGFA* gene, the ECs produced less (A) VEGFA and (K) CCL3 protein when co-cultured with *VEGFA*-silenced primary human IVD cells. The ECs also produced more (B) VEGFC and (J) PDGF-AA that are both positive regulators of lymphangiogenesis; MMPs (C-F; MMP1, MMP3, MMP10, MMP13) involved in matrix remodeling; chemokines including (G) G-CSF, (H) GM-CSF, (I) CXCL1, and (L) CCL5; and (J) PDGF-AA, a compensatory angiogenic growth factor. Units are pg/ml.

Taken together, this cross talk supports a mechanism in which endothelial cells, in the presence of IL-1β-stimulated secretome of VEGFA-silenced IVD cells, pivot toward a reparative state characterized by elevated lymphangiogenesis, increased immune cell engagement, and enhanced matrix remodeling—rather than the robust angiogenic response seen with IL-1β-stimulated VEGFA-intact IVD cells. These findings align with the vessel-pruning and tissue remodeling hypotheses seen in regenerative contexts and suggest that VEGFA inhibition not only reduces angiogenesis but promotes a more immunomodulatory and matrix-adaptive endothelial phenotype. In addition, several mediators known to support vessel homeostasis (e.g., Angiopoietin-2, Endoglin, Endothelin-1, PlGF) remained unchanged in co-culture with the VEGFA-silenced IVD cells (Table S2) and likely continue to support the homeostatic functions of existing vessels.

## DISCUSSION

This is the first study to demonstrate that intradiscal innervation contributes to low back pain behavior and locomotive performance in a model of intervertebral disc degeneration. While studies in the past have shown progressive associations between neurovascular structures with degeneration and pain^34–38^, none have been able to block or prevent these structures to prevent chronic pain and maintain locomotive performance. From this work, we also demonstrate that VEGFA may be a therapeutic target for a disease modifying therapy of low back pain. VEGFA ablation had a profound effect on neural and vascular infiltration and the subsequent mechanical sensitivity and locomotive performance following degeneration, independently of the structural and mechanical properties of the IVDs.

VEGFA plays a crucial role in limiting nerve and vessel infiltration post-injury. IVD degeneration in VEGFA-null mice exhibited reduced neurite and vessel lengths and the depths of their penetration, particularly in the outer annulus fibrosus (AF). VEGFA ablation inhibited the penetration and elongation of neural and vascular structures in the normally avascular IVD, preventing nociceptive sensory neurons from entering injured areas. The synergistic relationship between nerve and vessel infiltration highlights VEGFA’s importance in neurovascular coupling within the IVD, with its absence leading to diminished neurite and vessel growth, which may limit pathological nociceptive signals that propagate into chronic pain.

The loss of VEGFA also affected neuropeptide-related expression in the Dorsal Root Ganglion (DRG). The increase expression of TrKA, TRPV1, and TRPA1 in the DRG have been observed in chronic low back pain^39,40^, as well as in DRG that are sensitive to capsaicin stimulation^41^. VEGFA removal also decreased TRPA1 expression, whose expression is associated with nociception, in the DRGs. In contrast, TRPV1 expression, commonly associated with heat sensitivity^42^, did not change between VEGFA-null mice and wild-type mice. Indeed, the VEGFA-null mice and wild-type mice did not differ in heat sensitivity to the hot plate. VEGFA-null mice recovered from the early mechanical sensitivity at three-weeks and avoided the chronic sensitivity observed in WT mice. VEGFA-null mice did not develop the motor performance impairments exhibited by WT mice at later points (9-12 weeks), indicating a protective effect of VEGFA ablation against chronic pain and impaired performance. The asymmetric, injury-induced hypersensitivity was avoided by VEGFA inhibition, indicating that VEGFA loss mitigated the effects of IVD injury without altering overall pain sensitivity. Together, these findings suggest that VEGFA modulation is disease modifying rather than producing analgesic effects.

### Neurovascular Elongation vs Sprouting

Injuries to the intervertebral disc primarily trigger nerve and blood vessel sprouting as an adaptive response to inflammation and hypoxia^43,44^. This sprouting—stimulated by factors like NGF and inflammatory cytokines—leads to increased neurovascular infiltration into normally aneural and avascular disc regions, contributing to chronic pain and ongoing inflammation^45–47^. Consistent with this, we observed neurites and vessels infiltrating into the degenerating IVD at 3-weeks following injury. However, once sprouted, the neurites did not appear to elongate further, as evidenced by the relatively similar lengths and depths between 3- and 12- weeks. In this animal model, VEGFA is ubiquitously ablated from the whole animal, and it is not possible to determine the source of the VEGFA. However, our *in vitro* experiments support that blocking IVD-produced VEGFA is sufficient to prevent the growth of neurites and the formation of blood vessels. RNAseq analysis suggested that the reduced neural outgrowth is driven by the suppression of cell proliferation. We observed a phenotypic shift in the endothelial cells co-cultured with VEGFA-silenced IL-1β−stimulated IVD cells that secreted higher levels of VEGFC and PDGF-AA, positive regulators of lymphangiogenesis, alongside increased MMPs, and immune cell recruiting chemokines. Such changes suggest that, in the absence of VEGFA-driven angiogenesis, endothelial cells promote a microenvironment conducive to tissue remodeling and immune-mediated repair. These immune cell recruiting chemokines are likely to be important for the intrinsic healing of the IVD^48^. RNAseq analyses of the HMEC-1 cells confirms the ontological shift to lymphangiogenesis and matrix remodeling with a suppression of key angiogenic genes including *Heg1, Narap*, and *Ncs1*. There is also a suppression of endothelial cell-derived neurogenic factors including *Brsk2* and *Crlf1*.

Needle stab injury models in animals are frequently utilized to study the pathophysiology of intervertebral disc (IVD) degeneration and to develop treatments for chronic lower back pain (CLBP). These models effectively replicate critical aspects of human disc degeneration, such as mechanical damage, inflammatory responses, and biochemical changes. For example, they simulate the mechanical damage caused by traumatic events and the inflammatory cascade observed in human discs, which are crucial for understanding the progression of disc degeneration^49^. However, these models have limitations, especially in replicating the chronic, multifactorial nature of human CLBP and the complexity of human pain perception^50,51^. Human IVD degeneration occurs over years, with the herniation of the IVD being one of the most apparent instigating factors. Despite the varied etiologies of discogenic pain, the chronic symptoms and eventual degenerative presentation, such as the deterioration of IVD structure, are quite similar. Therefore, the needle injury model in murine IVD is employed to understand the mechanisms following IVD injury that eventually lead to low back pain symptoms. The needle injury in a mouse induces progressive, but relatively rapid degeneration, occurring over weeks and months, compared to human degeneration, which spans years and decades. Despite being traumatic relative to the human scale, this model is a necessary caveat for developing feasible and mechanistic experiments^51^. A single level lumbar injury utilized here may also underestimate the magnitude of our findings, since three-level injuries appear to show more pronounced effects^52^. Even with a single-level injury, however, the shift to painful behavior were obvious with longitudinal measurements. Notably, neurovascular ingrowth was more pronounced on the dorsal side opposite the ventral injury location,^34^, likely due to its proximity to the spinal cord and increased availability of neural and vascular sources. These findings have translational promise since it is possible to intervene early upon an acute IVD injury (e.g. a herniation) in the sequalae of chronic low back pain development. Since patients typically seek treatment only after an injury or an acute episode of pain, the timely post-injury inhibition of VEGFA may be able to prevent the transition to chronic low back pain.

Overall, the loss of VEGFA protected against mechanical sensitivity and locomotive performance and decreased intradiscal innervation. These findings show a causative relationship between neurovascular pathoanatomy and pain, while demonstrating VEGFA as a critical mediator of chronic pain development through the modulation of neurovascular features.

## MATERIALS AND METHODS

### Experimental Design

#### The Postnatal Inducible Ablation of Vegfa in Adult Mice

Young adult, 4–6-month-old, ubiquitin (UBC)-Cre^ERT2^ ; *Vegfa*^fl/fl^; Ai9-tdTomato^53^ mice on a C57BL/6J background^54–56^ allowed for a temporally controllable ablation of *Vegfa* in all tissues upon tamoxifen administration, along with an Ai9 fluorescence reporter to confirm recombination. The *Cre* element is activated when a mutant estrogen receptor (ERT2) renders *Cre* active upon tamoxifen binding and promotes Cre entering the nucleus to exise the *Vegf*a-encoding regions flanked by the *LoxP* sites. This approach avoids VEGFA-deficiency-mediated developmental defects and embryonic lethality, especially in pathways that are essential to organogenesis. All procedures (breeding, drug administration, procedures, and tissue extraction) were performed with WUSM IACUC (Institutional Animal Care and Use Committee) approval.

#### The Lumbar Puncture Intervertebral Disc Injury Model

Male and female mice of 4-6 months of age were randomly allocated into respective experimental groups. The animals undergoing surgery were anesthetized, shaven, and sterilized prior to surgery. Under microscopy guidance, the lateral lumbar spine was surgically exposed by mobilizing the peritoneum and retracting the psoas anteriorly. Using the superior pelvic margin to identify the L6 vertebrae, and then the L5/6 and L6/S1 IVDs are surgically exposed. The L5/6 IVD was injured using a 30G beveled hypodermic needle with three unilateral annular injuries. The sham animals received the same surgical exposure with no injury^20,22,34^.

On Days 3–4 following the surgery, Cre+;*Vegfa*^fl/fl^ and Cre−;*Vegfa*^fl/fl^ littermates received tamoxifen by gavage to induce *Vegfa* ablation in the Cre+ animals. (aka ‘VEGFA-null’) with Cre−;Vegfa^fl/fl^ littermates as WT animals. Data were collected at baseline, within 7 days, and at 3, 6, 9, and 12 weeks (behavioral assays); body weight at baseline/3/12 weeks; survival tracked for Kaplan–Meier analysis. Acute 3-week post-op and chronic 12-week post-op time points were chosen with the acute injury response allowing for the initial surgical wound healing recovery, while chronic LBP is defined as pain lasting 12 weeks or longer^57,58^. Additional experimental details are provided in the Supplemental Materials.

### Histological analyses

#### Sample Preparation

Spines from the euthanized animals were fixed in 4% paraformaldehyde, demineralized in 14% EDTA, and serially infiltrated with sucrose solutions prior to optimal cutting temperature (OCT) infiltration and freezing for cryosectioning.

#### Histopathological Evaluation of Degeneration

Histopathological scoring for degeneration in four compartments of the IVD including the NP, AF, end plate, and interface/boundary regions on 10μm-thick sections^59^. Mean degeneration scores from three independent investigators (n=3/sex/genotype/surgery, N=24 total) were used for analysis.

#### Structural Quantification of Nerve and Vascular Features by Immunohistochemistry

Two to three 50µm-thick serial sections were used to evaluate the neuronal and vascular morphology in 3-dimensions. Protein Gene Product 9.5 (PGP9.5), Endomucin, and DAPI counter stain^60^ were co-stained with respective antibodies to evaluate the nerve and vascular structures. DAPI, PGP9.5, and Endomucin helped visualize, respectively, cell nuclei, broad neural populations, and endothelial cells of vasculature via a confocal microscope at 2.5 μm focal depth increments with 10x magnification. Excitation and emission wavelengths were set as follows: DAPI excitation 405nm and emission 414-450 nm, PGP9.5 excitation 488nm and emission 498-530 nm, and Endomucin excitation 635nm and emission 650-725 nm. Neurite and vessel quantification of the outer AF were performed on max projections of a Z-stack with Fiji (ImageJ), with lengths on or within the AF and the surrounding fibrous tissue calculated from the Neuroanatomy SNT plugin and with depths calculated normal to the peripheral edge of the AF. PGP9.5 staining showed some non-specific regions so was only not included if neuronal morphology was apparent. To quantify spatial coupling of neurites and vessels, an automated MATLAB script was developed to measure lengths and depths and calculate colocalization lengths and a colocalization ratio based on distance from the rare neurite features, over a 30 µm distance. We also confirmed that 94% of *de novo* PGP9.5+ neurites that infiltrate the IVD are positive for Na_v_1.8, a sensory neuron marker (Figure S3).

#### Neuropeptide and Ion Channel Immunostaining of the Dorsal Root Ganglia

Frozen sections of isolated Doral Roo Ganglia (DRG) from L1-L3 levels were processed for immunohistochemistry for anti-TrkA, anti-TRPA1, and anti-TRPV1. Nuclear counterstaining was performed using DAPI, and slides were mounted prior to imaging. DRG immunohistochemical images were quantified using the custom Python software, DRGVisual. For each cell, the corrected total cell fluorescence (CTCF) was calculated by subtracting the background intensity from each pixel’s intensity within the cell region and summing the total fluorescence across the cells for each ion channel.

### 3D Structural Evaluation of Intervertebral Disc Structure Using Contrast-enhanced µCT

At 3 weeks post lumbar intervertebral disc injury, *in vivo* µCT was used to evaluate structural changes of the intradiscal space using disc height^20^. The structural changes at this 3 week post injury time point were further evaluated using the e*x vivo* contrast-enhanced micro-computed tomography (CEμCT). Ioversol was used as a contrast agent to highlight the water-rich nucleus pulposus^61^. Total disc ROI and nucleus pulposus ROI were contoured using the MATLAB graphical user interface from Washington University Musculoskeletal Image Analyses (github.com). NP hydration, a ratio of nucleus attenuation to total disc attenuation (NI/DI), was calculated along with volumes and disc height ratio, DHR, as an average of 5 height measures in a midsagittal slice over the disc width^62^.

### Biomechanical Assessment of the Intervertebral Disc

Mechanical behavior of the IVDs using dynamic compression testing under displacement control^63^. FSUs were affixed to aluminum platens and immersed in a phosphate-buffered saline bath at room temperature and preloaded to 0.25N. A sinusoidal compressive waveform at 10% strain and 1 Hz frequency was applied for 20 cycles. The average stiffness was calculated from the second through the final loading cycles and dissipation factor (aka the loss tangent), was calculated from the phase shift between load and displacement data.

### Behavioral Assessments

Habituation measures were taken to reduce animal stress during behavioral assessments^64^. All tests were performed by the same laboratory personnel who was blinded to genotype and surgical procedure. Baseline data was collected one week prior to surgery and post-operative data collection included a cross-sectional group at baseline, 3, and 12 weeks and a longitudinal group at 3, 6, 9, and 12 weeks.

#### Mechanical Sensitivity Measured Using the Von Frey and E-Von Frey

Mechanical sensitivity was measured using the up-down method of the Von Frey test using Semmes Weinstein filaments and an electronic Von Frey (EVF; BIOSEB, France)^65,66^. Prior to baseline behavioral testing, mice were habituated to the elevated mesh grid chamber for 1 hr. Calibrated filaments were applied to the plantar surface of the hind paw to elicit paw withdrawal or filament buckling^67^. Filament Von Frey was done on the left hind paw to focus on the injury response. EVF consisted of maximum force values at withdrawal from the median of the result of 3 trials which were conducted bilaterally, with at least 5 min between testing opposite paws, and at least 10 min between testing the same paw. Post-op measures were normalized to the pre-surgery baseline.

#### Hot and Cold Nociceptive Sensitivity

Mice were subjected to thermal stimulus to measure the temperature sensitivity response^68,69^. A hot/cold plate (Bioseb, France) was set to 55±0.5°C or 5±0.5°C and mice were placed individually onto the plate with a transparent enclosure to prevent escape and allow for visibility. The latency to the first sign of nociceptive behavior, either lick of the hind paws or jumping for hot plate; and lifting, licking, or shaking of the hind paws for cold plate was recorded with a cutoff time of 20 seconds for hot or 30 seconds for cold to prevent tissue damage. Three repeated measures were collected with >15 minutes between tests to allow for home cage recovery.

### Functional Assessments of Performance

#### Locomotor Function

##### Rotarod

Assessing motor coordination and balance, the Rotarod performance test challenged mice to a rotating rod apparatus (Bioseb, France)^66,70,71^. Prior to testing, mice were trained on the Rotarod at a slow 4 rpm for 2 minutes to ensure baseline capabilities. On a subsequent day, mice were tested on an accelerating bar from 4 to 40 rpm over a 2-minute period. Latency to fall was recorded for each mouse. Each mouse underwent 5 trials with a >5-minute rest period between trials to reduce fatigue. The best performance of the 5 trials was used for data analysis and normalized to baseline.

##### Open Field Testing

In the open field test, designed to evaluate exploratory passive behavior, mice were individually introduced into the center of an open field arena (40 cm x 40 cm, with 30 cm high walls) equipped with an automated tracking system (Omnitech Electronics, Columbus, OH)^66,70,71^. Total distance traveled and instances of vertical rearing were recorded for 60 minutes.

#### Strength

##### Inverted Wire Hang Endurance

In the wire hang behavioral test, mice were evaluated for strength and endurance^72^. The apparatus consisted of a wire mesh (2 mm diameter) suspended above a soft bedding layer to ensure a safe landing for the mice. Each mouse was gently held by the base of the tail and allowed to grasp the wire at a 60° incline before the mesh was inverted to 180°. The duration the mouse remained suspended inverted was recorded up to a maximum of 2 minutes. A minimum 5-minute rest was given between trials. The maximum time of three trials was recorded.

##### Grip Strength

Grip strength was measured using a uniaxial grip force tester (Bioseb, France)^73^. Hind paw grip strength assessed mechanical strength as the mouse was gently restrained and allowed to freely grab and hold a bar with both hind paws. Maximum force was recorded as the tail was pulled in a steady manner to avoid rate-dependent effects. A brief rest period, >15 sec, was given between each of the 5 repeated trials. Average strength was reported normalized to body weight at the time of testing and normalized to pre-surgery baseline.

### Cell Culture

#### Human Primary IVD Cell Culture

Primary human intervertebral disc cells were isolated from the disc tissue of patients undergoing elective discectomy surgeries (Table S3). Human biospecimens were obtained under IRB Exemption Category 4, in accordance with institutional and federal guidelines for research using de-identified, previously collected samples^74,75^. Isolated cells were centrifuged, resuspended, plated, and incubated at 37°C, 5% CO^2^, 20% O^2^ to grow to 90% confluence.

#### Vascular Endothelial Growth Factor (VEGF)-siRNA Transfection

Isolated human IVD cells were seeded in 6-well plates at a density of 3×10^5^ cells per well 24 hours prior to transfection. VEGFA-specific siRNA (sc-29620; Santa Cruz Biotechnology, Dallas, Tx) and scrambled control siRNA-FITC (sc-36869; Santa Cruz Biotechnology) were transfected into IVD cells using Lipofectamine™ RNAiMAX Transfection Reagent (Catalog #: 13778100, Thermo Fisher) following the manufacturer’s protocol. The final concentration of siRNA in the culture medium was 50 pM. The effectiveness of transfection was evaluated by checking the amount of secreted VEGFA in media using ELISA (DY293; R&D system, Minneapolis, MN) and confirming transfection of control siRNA by FITC fluorescence with a Nikon Eclipse Ti2 inverted scope at 10x.

#### Co-culturing VEGFA-silenced IVD cells with Immortalized Human Microvascular Endothelial Cell (HMEC-1) and Neuroblastoma Cell (SH-SY5Y) cultures

IVD cells were plated in a 6 well plate and transfected with either control, scrambled siRNA or with VEGFA siRNA. Medium with 1% FBS and recombinant human interleukin-1β (IL-1β; R&D Systems) at 1 ng/mL was added to stimulate Scram siRNA (IL-1β+Scram siRNA) and VEGFA siRNA (IL-1β+VEGFA siRNA) transfected IVD cells for 24 hours. After stimulation, HMEC-1 cells “ECs” that were plated for 24 hours on 1 µm pore co-culture inserts (Thincert cell culture insert; Greiner, Monroe, NC) at a density of 3×10^5^ cells were inserted to the wells containing Scram siRNA or VEGFA siRNA transfected IVD cells with MCDB media plus 1% FBS.

HMEC-1 (aka Endothelial cells - ECs) and SH-SY5Y (aka Neuronal cells - NCs) cells were obtained from the suppliers, cultured and passaged 5-10 times, and then grown to 90% confluence. ECs co-cultured with stimulated Scram siRNA treated IVD cells are denoted as EC_IL-1β+Scram siRNA_ and ECs co-cultured with stimulated VEGFA siRNA treated IVD cells are denoted as EC_IL-1β+VEGF siRNA._ Naïve ECs that did not undergo co-culturing were used as a control group. Following coculture, the ECs underwent a Matrigel tube formation assay^76^, RNA-sequencing, and a multiplex ELISA of the EC-derived conditioned media (Figure S4). NCs were plated at a density of 1.5 × 10^5^ cells per well in a 6 well plate and cultured for 48 hours prior to the co-culture experiment. For co-culturing, non-stimulated IVD cells “naive”, IL-1β+Scram siRNA, and IL-1β+VEGFA siRNA transfected IVD cells were plated on the co-culture insert at a density of 3.0 × 10^5^ cells and co-cultured for 48 hours with NCs. Following co-culture, the NCs underwent a neurite outgrowth assay and RNA-sequencing (Figure S5).

#### RNA Sequencing and Multiplex protein ELISA

Post co-culture experiments, ECs and NCs were lysed, the total RNA was extracted and sequenced on an Illumina NovaSeq X Plus. RNA-seq data were processed using standard pipelines for base calling, alignment, quantification, and normalization. Quality control ensured accurate read alignment, expression quantification, and removal of low-quality or low-expression genes. Normalized expression values were then used for downstream differential analysis. The raw data files are available on the Gene Expression Omnibus database: GSE294577. The conditioned media from naive ECs and those co-cultured with IL-1β+Scram siRNA and IL-1β+VEGFA siRNA transfected IVD cells was collected and analyzed by multiplex protein assay.

### Statistical and Bioinformatics Analyses

Statistical comparisons on repeated measures, eg the longitudinal behavioral assays, were done using a linear mixed effect model to assess surgery (Sham/Injured), genotype (WT/VEGFA-null), 3- and 12-weeks after surgery (3/12), sex (M/F), and surgery (Naïve/Sham). Linear mixed effect models were initiated with all factors and reduced in complexity as factors were found to be not significant. Additionally, focused questions were modeled to assess injured groups with genotype and time. A 2 or 3 factor ANOVA was used to assess effects of injury, genotype, and sex on CEµCT, biomechanics, histopathologic and immunohistochemical outcomes. ANOVA pairwise comparisons were evaluated with the Fisher’s Least Significant Difference method. Survival curves implemented Mantel-Cox test while binary penetration results used a Fisher’s exact test. Simple pairwise comparisons were done using a Welch’s t-test when appropriate. Appropriate adjustments for unequal variances and non-normal distributions were made where appropriate. The ROUT outlier test with Q=0.1% removed definitive outliers within groups^77^. R and GraphPad Prism were used to model the results. For gene ontology analyses, biological pathways enriched by the upregulated and downregulated DEGs were identified by using the Panther Classification System with version Panther 19.0 by using the statistical overrepresentation test. For ECs, an adjusted p value of p < 0.05 and logFC of 0.41 was used to identify statistically significant DEGs with at least a 50% increase in fold change of the treatment group when compared to the control. For neuronal cells, an adjusted p value of p < 0.05 and logFC of 0.41 was used to identify statistically significant DEGs with at least a 50% increase in fold change of the treatment group when compared to the controls. The full list of all DEGs and GO terms for the endothelial cell’s comparisons are listed in Tables S8A and S8B and for the neuronal cell comparisons are listed in Tables S9A and S9B. Additional details on experimental methods can be found in the Supplemental Materials.

## SUPPLEMENTARY MATERIALS

Supplementary Materials.docx

DEGs Endothelial Cells.xlsx

DEGs Neuronal Cells.xlsx

Gene Ontology Neuronal Cells.xlsx

Gene Ontology Endothelial Cells.xlsx

## List of Abbreviations

AF: Annulus Fibrosus
Ai9: Molecular reporter expressing tdTomato fluorescent protein (Rosa-CAG-LSL-tdTomato)
ANOVA: Analysis of Variance
CEµCT: Contrast-Enhanced Micro-Computed Tomography
cLBP: Chronic Low Back Pain
CM: Conditioned Media
CTCF: Corrected Total Cell Fluorescence
C57BL/6J: Common inbred mouse strain
DAPI: 4′,6-Diamidino-2-Phenylindole (DNA stain)
DEG: Differentially Expressed Gene
DHR: Disc Height Ratio
DRG: Dorsal Root Ganglion
EC: Endothelial Cell (HMEC-1)
EDTA: Ethylenediaminetetraacetic Acid (chelating agent)
ELISA: Enzyme-Linked Immunosorbent Assay
EMCN: Endomucin (endothelial cell marker)
EMEM, ATCC: Eagle’s Minimum Essential Medium, from American Type Culture Collection
ERT2: Modified Estrogen Receptor Ligand Binding Domain
EVF: Electronic Von Frey
FBS: Fetal Bovine Serum
FITC: Fluorescein Isothiocyanate dye
FSU: Functional Spinal Unit
HBSS: Hanks’ Balanced Salt Solution
HMEC-1: Human Microvascular Endothelial Cell Line
IL1β: Interleukin 1 Beta
IVD: Intervertebral Disc
LBP: Low Back Pain
LoxP: Locus of X-over P1 (recombination sites for Cre-Lox systems)
LSD: Least Significant Difference (statistical post hoc test)
MCDB: Molecular, Cell, and Developmental Biology
Nav1.8: Voltage-Gated Sodium Channel (gene SCN10A) expressed in nociceptive neurons
NC: Neuron cells (SH-SY5Y)
NFkB: Nuclear Factor Kappa B
NGF: Nerve Growth Factor
NI/DI: Nucleus Attenuation Intensity/Disc Attenuation Intensity
NP: Nucleus Pulposus
OCT: Optimal Cutting Temperature Compound
PAP: Hydrophobic barrier pen
PBS: Phosphate-Buffered Saline
PGP9.5: Protein Gene Product 9.5 (neuronal marker)
P/S: Penicillin/Streptomycin
ROI: Region of Interest
SCRAM: Scrambled siRNA (negative control)
SH-SY5Y: Human Neuroblastoma Cell Line
siRNA: Small Interfering RNA
SNT: Simple Neurite Tracer (ImageJ plugin)
TNFα: Tumor Necrosis Factor Alpha
TNT: Tris-NaCl-Tween buffer
TrKA: Tropomyosin Receptor Kinase A (NGF receptor)
TRPA1: Transient Receptor Potential Ankyrin 1
TRPV1: Transient Receptor Potential Vanilloid 1
UBC: Ubiquitin
VEGFA: Vascular Endothelial Growth Factor Type A
WT: Wild Type

## Acknowledgments

We acknowledge the support of the Washington University Musculoskeletal Research Center Structure and Function Core (Michael Brodt); Histology Core (Crystal Idleburg and Samantha Coleman); and Animal Behavior and Function Core. We also acknowledge Kelsey Collins and Washington University Pain Center Judith Golden for their training in behavioral assessments of pain. We thank the Washington University Genomics Technology Access Core (GTAC), and the Eve-tech for the multiplex ELISA analysis.

## Author contributions

Conceptualization: RSP, MJS, LAS, MCG, SYT

Methodology: RSP, HJM, SWC, REW, ALL, ANS, LJ, EGB, JAM, ATB, LAS, MCG, SYT

Data Interpretation: Methodology: RSP, REW, HJM, ALL, ANS, SWC, LJ, LAS, MCG, SYT

Investigation: RSP, HJM, SWC, ALL, LJ, LAS, MCG, SYT

Visualization: RSP, HJM, SWC, ALL, SWC Funding acquisition: LAS, MCG, SYT

Project administration: RSP, SYT Supervision: SYT

Writing original draft: RSP, HJM, SWC, ALL

Writing review & editing: RSP, SWC, SYT

## Competing interests

Provisional Patent: 020859/US

Title: VEGFA INHIBITORS AS A TREATMENT FOR CHRONIC MECHANICAL ALLODYNIA

Co-inventors: RSP, HJM, MCG, SYT

## Data and materials availability

All data, code, and materials used will be made available to other researchers upon reasonable request for the proposes of reproducing or extending the analysis. Vegfa^fl/fl^ mice were obtained via a materials transfer agreements (MTAs) with Genentech Inc and provided by MJS. The raw data files of the endothelial cell and neuron gene expression analyses are available on the Gene Expression Omnibus database under accession number GSE294577. Supplemental materials contains additional methods and data in support of this study.

