## Supplemental Materials for "VEGFA critically controls neurovascular invasion and chronic low back pain during intervertebral disc degeneration"

Supplemental Methods

*Confirmation and Quantification of Vegfa Ablation*

The Ai9-tdTomato reporter protein is co-expressed with Cre recombinase presence and is visibly detectable (Figure S1) after two doses of tamoxifen (100 mg/kg on two consecutive days). We also evaluated the functional efficiency of *Vegfa* ablation in the intervertebral disc (IVD) by measuring the production of the VEGFA protein secreted by the IVD. Mice with Cre- and Cre+ alleles driven by the ubiquitin promoter in animals with the homozygous *Vegfa* floxed alleles (UBC-Cre$^{ERT2}$; *Vegfa*$^{fl/fl}$ Ai9) received 2 consecutive days of tamoxifen administration via oral gavage (100 mg/kg in corn oil). The animals were sacrificed on the third day, following the two days of induction. The functional spinal units, FSUs (vertebra–intervertebral disc–vertebra units), were cultured as previously described and stimulated with TNF-α (5 ng/ml) to provoke an inflammatory response[1]. Media were collected every other day for 7 days and analyzed using a VEGFA ELISA (R&D Systems Cat: DY493).

*Injury-induced Intervertebral Disc Degeneration*

The animals undergoing surgery were first anesthetized using subcutaneous lidocaine injection (0.5% at 7mg/kg) and sustained by isoflurane inhalation (3–4% v/v induction, 2–2.5% v/v maintenance at 1 L/min). The left flank was shaved and sterilized (70% ethanol, povidone–iodine). Under microscopic guidance, a retroperitoneal dissection was performed to expose the lateral aspect of the spine using a Penfield dissector and size 11 scalpels. The pelvis and hip were rotated posteriorly to enhance the working space. The peritoneal wall was gently tugged to expose the psoas muscle, which was retracted posteriorly to anteriorly using a cotton swab and then held in place with a metal spatula. The spinal column and IVDs at the L5/L6 and L6/S1 levels were exposed by blunt dissection. The superior margin of the pelvic bone indicated the L6 vertebral body, confirming the locations of L5/L6 and L6/S1 IVDs. The L5/L6 IVD was targeted and either exposed in shams or injured with a 30G needle 3 times through the annulus fibrosus and partially into the nucleus pulposus. Postoperatively, mice were monitored every 8–12 hours for 4 days. Pain management included intraperitoneal injections of carprofen (5 mg/kg) every 8–12 hours, supplemented with carprofen tablets (2 mg/tablet) for 48-72 hours.

*Tamoxifen administration to induce genetic recombination*

On days 3 and 4 postoperatively, tamoxifen (TAM, Cat#: T5648, Millipore Sigma) was given via oral gavage (100 mg/kg in corn oil, Cat#: C8267, Millipore Sigma) to Cre+;*Vegfa*$^{fl/fl}$ animals ('VEGFA-null') and Cre-;*Vegfa*$^{fl/fl}$ littermates ('WT') inducing *Vegfa* ablation. Following the two-day tamoxifen administration, recombination can be visually observed in the Cre+ animals (Figure S1A).

**Histological analyses**

*Sample Preparation*

Spines from the euthanized animals were fixed in 4% paraformaldehyde for approximately 48 hours, washed 6 times for 15 minutes each in 1X PBS, and subsequently demineralized in 14% EDTA for 10-14 days. After which, samples were washed 8-10 times for 15 minutes in 1X PBS and serially infiltrated with 10%, 20%, and 30% sucrose solutions for 1 hour each prior to optimal cutting temperature (OCT) infiltration and freezing for cryosectioning.

*Histopathological Evaluation of Degeneration*

Midsagittal 10-μm thick sections were stained with Safranin O/Fast Green and imaged at 10× on a Nanozoomer (Hamamatsu, Japan). Degeneration was scored in the NP, AF, end plate, and interface/boundary regions[2]. Mean degeneration scores from three independent investigators (n = 3/sex/genotype/surgery, N = 24 total) were used for analysis.

*Structural Quantification of Nerve and Vascular Features by Immunohistochemistry*

Sections 50-μm-thick were used to evaluate the neuronal and vascular morphology in 3-dimensions. Serial sections were collected with CryoJane adhesive slides to adhere tissue to the slide during IHC staining with tissues being stored at -80°C prior to antibody staining.

To stain for Protein Gene Product 9.5 (PGP9.5), Endomucin, and DAPI counter stain, a 4-day protocol[3] was followed whereby slides were dried at room temperature for 30 minutes and washed in PBS 2 times for 5 minutes. A PAP pen was used to create a hydrophobic perimeter around the tissue and allow for direct pipetting of reagents onto the slides to avoid tissue loss from vertical slide immersion. Slides were blocked with 10% donkey serum and 0.3% Triton X-100 in PBS for 1 hour at room temperature. The primary PGP9.5 antibody (Millipore Cat # Ab1761) at a 1:1000 dilution and Endomucin antibody (Thermo Fisher/eBioscience, Cat #14-5851-85) at a 1:500 dilution were added to 1% normal donkey serum in a TNT (Tris-NaCl-Tween) buffer and incubated for 48 hours at 4°C in a humidified chamber. Samples were then washed 3 times for 5 minutes in TNT buffer prior to incubation of a PGP9.5 secondary antibody (Jackson, Cat #711-545-152) at 1:1000 concentration and 488-nm fluorophore and Endomucin secondary antibody (Jackson Cat #712-605-153) at 1:1000 concentration and 657-nm fluorophore in 1% normal donkey serum in TNT buffer for 24 hours at 4°C in a humidified chamber. Samples were washed 3 times with TNT for 5 minutes prior to DAPI (Sigma-Aldrich D9542) at 1:1000 for 5 minutes at room temperature. A final wash with TNT 3 times for 5 minutes was done before cover slipping and sealing with Fluoromount-G the next day (Thermo Cat:00-4958-02).

DAPI, PGP9.5, and Endomucin helped visualize, respectively, cell nuclei, broad neural populations, and endothelial cells of vasculature via a confocal microscope (Leica, USA) at 2.5 μm focal depth increments with 10× magnification and 2048×2048 resolution and 600 speed with 2 frame averaging. Excitation and emission wavelengths were set as follows: DAPI excitation 405 nm and emission 414-450 nm, PGP9.5 excitation 488 nm and emission 498-530 nm, and Endomucin excitation 635 nm and emission 650-725 nm. There was no spectral overlap with Ai9 tdTomato (excitation 554 nm, emission 581 nm). Neurite and vessel quantification of the outer AF was performed on maximum-intensity projections of a Z-stack with Fiji (ImageJ), with lengths on or within the AF and the surrounding fibrous tissue calculated from the Neuroanatomy SNT plugin and with depths calculated normal to the peripheral edge of the AF. Because PGP9.5 showed some nonspecific staining, only regions with clear neuronal morphology were included. To confirm that the PGP9.5 positive structures within the IVD accurately captured the sensory-neuron population, we conducted a direct comparison of the PGP9.5 staining of an injured intervertebral disc in a $Na_v1.8$ td Tomato reporter mouse and found the $Na_v1.8+$ features accounted for 88% of the $PGP9.5^+$ features with spatial colocalization (R = 0.93, Figure S3). The sodium-sensitive ion channel $Na_v1.8$ is highly specific to sensory neurons[4-6].

To quantify spatial coupling of neurites and vessels, an automated MATLAB script was developed to measure lengths and depths and calculate colocalization lengths and a colocalization ratio based on distance from the rarer neurite features. Spatial coupling was defined as vessels within 30 μm from neurites and based on the distance between two adjacent lamellae because lamellar structure.

*Neuropeptide Immunostaining of the Dorsal Root Ganglia*

Frozen DRG sections, isolated from L1-L3 were processed for immunohistochemistry using a standardized protocol across all targets. Briefly, slides were fixed in 4% paraformaldehyde for 10 minutes at room temperature. After washing in 0.1 M Tris buffer (pH 7.6), nonspecific binding was minimized through sequential blocking steps with 0.005% BSA in Tris and 10% normal goat serum. Sections were incubated overnight at 4°C with primary antibodies—anti-TrkA (1:100, Alomone Labs #ANT-018), anti-TRPA1 (1:200, Alomone Labs #ACC-037), or anti-TRPV1 (1:500, Abcam #ab6166)—diluted in Tris-BSA. After rinsing, a goat anti-rabbit Alexa Fluor 488 secondary antibody (1:300, Invitrogen #A11006) was applied for 4 hours at room temperature or overnight at 4°C. Nuclear counterstaining was performed using DAPI (1:5 dilution in 0.1 M Tris), and slides were mounted prior to imaging. DRG immunohistochemical images were quantified using the custom Python software, DRGVisual. Contrast Limited Adaptive Histogram Equalization (CLAHE) was applied to enhance the contrast of the images. The processing began with Canny edge detection to refine the segmentation and ensure accurate depiction of cell boundaries. Following edge detection, seed points within the cells were identified to indicate the center of the cells, and diameters were drawn to estimate each cell's maximum size. Cell regions were grown using a projection-based region-growing algorithm that expands the region from each seed point by examining neighboring pixels and including them in the region if they meet criteria based on intensity and connectivity. For each cell, the corrected total cell fluorescence (CTCF) was calculated as the sum of background-subtracted pixel intensities within each cell for each ion channel.

#### 3D Structural Evaluation of Intervertebral Disc Structure Using Contrast-enhanced μCT

At 3 weeks post lumbar intervertebral disc injury, in vivo μCT was used to evaluate structural changes of the intradiscal space (VivaCT, 45 kVp, 177 μA, 300 ms integration time, 10 μm voxel) with isoflurane inhalation (2-3% v/v). Disc height was calculated in ImageJ as the average of 5 measures across the midsagittal view[7]. To confirm that the loss of VEGFA did not affect the structure of the IVD, we examined the uninjured L4/L5 and L6/S1 IVDs between the VEGFA-null and the WT animals. Additionally, we examined the injured L5/L6 IVD level to ensure that the VEGFA-null animals were susceptible to injury-mediated degeneration.

The structural changes at this 3-week post injury time point were further evaluated using the e*x vivo* contrast-enhanced micro-computed tomography (CE-μCT). Ioversol was used as a contrast agent to highlight the water-rich nucleus pulposus[8]. The fresh IVDs were immersed in 50% Ioversol (OptiRay 350, Guerbet Pharma) at 37°C for 24 hours before imaging. The Scanco40 μCT system (Zurich, Switzerland) scanned samples at 45 kVp, 177 μA, 10.5 μm voxel size, and 300 ms integration time, after incubation[9]. Groups were n=3 mice/sex/genotype/injury. Total disc ROI and nucleus pulposus ROI were contoured using the MATLAB graphical user interface from Washington University Musculoskeletal Image Analyses (github.com). NP hydration, a ratio of nucleus attenuation to total disc attenuation (NI/DI), was calculated along with volumes and disc height ratio, DHR, as an average of 5 height measures in a midsagittal slice over the disc width.

#### Biomechanical Assessment of the Intervertebral Disc

Mechanical behavior of the IVDs using dynamic compression testing under displacement control was performed using a BioDent reference point indenter (Active Life Scientific)[10]. After CE-μCT, these samples (n = 3 mice per sex/genotype/injury) were mechanically tested with the strain rate calculated from the 5-point average disc height. FSUs were affixed to aluminum platens and immersed in a phosphate-buffered saline bath at room temperature and preloaded to 0.25 N. A sinusoidal compressive waveform at 10% strain and 1 Hz frequency was applied for 20 cycles. The average stiffness was calculated from the second through the final loading cycles and the dissipation factor (aka loss tangent) was calculated from the phase angle between load and displacement data.

**Behavioral Assessments**

Habituation measures were taken to reduce animal stress during behavioral assessments. Acclimation to room environment prior to assessment was set for 1 hour at the beginning of each behavioral testing week. To avoid additional animal stress, assessments were typically performed twice daily between 9:00 AM and 5:00 PM, with 1-hour rests between tests to minimize diurnal variations in activity[67]. All tests were performed by the same laboratory personnel. Baseline data were collected one week prior to surgery and postoperative data collection included a cross-sectional group at baseline, 3, and 12 weeks and a longitudinal cohort at 3, 6, 9, and 12 weeks. Care was taken to ensure a clean, quiet, and odor-free environment at a controlled temperature (20-23 °C). Testing was initiated up to one week prior to the 3- and 12-week time points as the full set of assays took 4-5 days to complete with efforts made to reproduce the order and timing of each test to remove stress on the animals and replicate methods.

*Mechanical Sensitivity Measured Using the Von Frey and E-Von Frey*

Mechanical sensitivity was measured using the up-down method of the Von Frey test using Semmes-Weinstein filaments (Bioseb, France) and an electronic Von Frey (EVF; BIOSEB, France) [11,12]. Prior to baseline behavioral testing, mice were habituated to the elevated mesh grid chamber for 1 hr. Calibrated filaments were applied to the plantar surface of the hind paw to elicit paw withdrawal or filament buckling[13]. Filament testing was performed on the left hind paw to focus on the injury response. EVF consisted of maximum force values at withdrawal from the median of the result of 3 trials which were conducted bilaterally, with at least 5 minutes between testing opposite paws, and at least 10 minutes between testing the same paw. Postoperative measures were normalized to baseline.

*Hot and Cold Nociceptive Sensitivity*

Mice were subjected to thermal stimulus to measure the temperature sensitivity response[14,15]. A hot/cold plate (Bioseb, France) was set to 55±0.5°C or 5±0.5°C and mice were placed individually onto the plate with a transparent enclosure to prevent escape and allow for visibility. The latency to the first nocifensive behavior seen in hind-paw licking or jumping (hot plate) and hind-paw lifting, licking, or shaking (cold plate) was recorded with a cutoff time of 20 seconds for hot or 30 seconds for cold to prevent tissue damage. Three repeated measures were collected with >15 minutes between tests to allow for home cage recovery. Values were averaged and normalized to baseline.

Functional Assessments of Performance

*Locomotor Function – Rotarod*

Assessing motor coordination and balance, the Rotarod performance test challenged mice to a rotating rod apparatus (Bioseb, France)[12,16,17]. Prior to testing, mice were trained on the Rotarod at a slow 4 rpm for 2 minutes to ensure baseline capabilities. Mice unable to achieve this baseline capacity were excluded from assessment. On a subsequent day, mice were tested on an accelerating rod (4 to 40 rpm over 2 minutes). Latency to fall, measured in seconds, was recorded for each mouse. Each mouse underwent 5 trials with a >5-minute rest period between trials to reduce fatigue. The best performance of the 5 trials was used for data analysis and normalized to baseline when possible.

*Open Field Testing*

In the open field test, designed to evaluate locomotor/exploratory behavior, mice were individually introduced into the center of an open field arena (40 cm x 40 cm, with 30 cm high walls) equipped with an automated tracking system (Omnitech Electronics, Columbus, OH)[12,16,17]. The apparatus featured an array of infrared sensors for precise movement tracking of total distance traveled and vertical rearing over a 60-minute test period.

Strength

*Inverted Wire Hang Endurance*

In the wire hang behavioral test, mice were evaluated for neuromuscular strength and endurance[18]. The apparatus consisted of a wire mesh (2 mm diameter) suspended above a soft bedding layer to ensure a safe landing for the mice. Each mouse was gently held by the base of the tail and allowed to grasp the wire at a 60° incline before the mesh was inverted to 180°. The duration the mouse remained suspended inverted was recorded up to a maximum of 2 minutes. A minimum 5-minute rest was given between trials. The maximum time of three trials was recorded.

*Grip Strength*

Grip strength was measured using a uniaxial grip force tester (Bioseb, France)[19]. Hind paw grip strength assessed mechanical strength as the mouse was gently restrained and allowed to freely grab and hold a bar with both hind paws. Maximum force was recorded as the tail was pulled in a steady manner to avoid rate-dependent effects. A brief rest period, >15 seconds, was given between each of the 5 repeated trials. Average strength was reported normalized to body weight at the time of testing and to baseline.

### Cell Culture

*Human Primary IVD Cell Culture*

Human intervertebral disc cells were isolated from the IVD tissue of eight patients (mean age ± SD = 68.5 ± 8.1; female:male = 5:3; mean Pfirrmann grade ± SD = 3.5 ± 0.7) removed during elective discectomy surgeries (Table S3). Human biospecimens were obtained under IRB Exemption Category 4, in accordance with institutional and federal guidelines for research using de-identified, previously collected samples, as described in prior studies[20,21]. Tissue specimens were placed in sterile Ham's F-12 medium (F-12; Gibco-BRL, Grand Island, NY) containing 1% penicillin/streptomycin (P/S; Gibco-BRL) and 5% fetal bovine serum (FBS; Gibco-BRL). Disc tissue was washed three times with Hank's balanced salt solution (HBSS; Gibco-BRL) containing 1% P/S to remove blood and other bodily contaminants prior to isolation. Specimens were minced and digested for 60 min at 37°C under gentle agitation in F-12 medium containing 1% P/S, 5%

FBS, and 0.2% pronase (Calbiochem, La Jolla, CA). The specimens were rinsed three times with HBSS and 1% P/S and incubated overnight in a solution of F-12 medium containing 1% P/S, 5% FBS, and 0.025% collagenase I (Sigma Chemical Co, St. Louis, MO). A sterile nylon mesh filter (70 µm pore size) was used to separate suspended cells from the remaining tissue debris. Isolated cells were centrifuged (2000 rpm, 5 min) and resuspended in F-12 medium with 10% FBS and 1% P/S. Cells were plated in a 75-$cm^2$ culture flask (VWR Scientific Products, Bridgeport, NJ) and incubated at 37°C, 5% $CO_2$, 20% $O_2$ in a humidified incubator. Culture medium was changed three times weekly, and cultures grew to 90% confluence. Cells were then trypsinized (0.05% trypsin/ethylene diaminetetraacetic acid; Gibco-BRL) and re-plated (6 x $10^5$ cells/mL) in 75-$cm^2$ culture flasks.

*Immortalized Human Microvascular Endothelial Cell (HMEC-1) Culture*

HMEC-1(CRL-3243; ATCC, Manassas, VA) were cultured in MCDB 131 medium (10372019; Gibco, Thermo Fisher Scientific) containing 10 ng/mL epidermal growth factor (PHG0311; Thermo Fisher Scientific, Waltham, MA), 500 µg/mL hydrocortisone (H0396; Sigma), 1% P/S, 2 mM L-Glutamine and supplemented with 10% FBS. Cells at passage number 5-10 were plated in 75-$cm^2$ culture flasks and grown to 90% confluence.

*Immortalized Human Neuroblastoma Cell (SH-SY5Y) Culture*

The SH-SY5Y human neuroblastoma cell line (CRL-2266; ATCC) was cultured in a 1:1 mixture of Eagle's Minimum Essential Medium (EMEM; ATCC, Manassas, VA) and Ham's F12 medium, supplemented with 10% FBS and 1% P/S in a 5% carbon dioxide humidified incubator at 37°C. Cells at passage number 5-10 were plated in 75 $cm^2$ culture flasks and grown to 90% confluence.

*Vascular Endothelial Growth Factor (VEGF)-siRNA Transfection*

Isolated human IVD cells were seeded in 6-well plates at a density of $3x10^5$ cells per well 24 hours prior to transfection. VEGF-specific siRNA (sc-29620; Santa Cruz Biotechnology, Dallas, TX) and scrambled control siRNA- FITC (sc-36869; Santa Cruz Biotechnology) were transfected into the IVD cells using Lipofectamine™ RNAiMAX Transfection Reagent (Catalog #: 13778100, Thermo Fisher) following the manufacturer's protocol. The final concentration of siRNA in the culture medium was 50 pM. The effectiveness of transfection was evaluated by checking the amount of the secreted VEGF in media using ELISA (DY293; R&D Systems, Minneapolis, MN) and confirming transfection of control siRNA by FITC fluorescence with a Nikon Eclipse Ti2 inverted scope at 10×.

*Co-culture between IVD cells and HMEC-1 (ECs) Cells*

IVD cells were plated in a 6 well plate and transfected with either control, scrambled siRNA (Scram siRNA) or with VEGF specific siRNA (VEGF siRNA). Medium with 1% FBS and recombinant human interleukin-1β (IL-1β; R&D Systems) at 1 ng/mL was added to stimulate Scram siRNA (IL-1β+Scram siRNA) and VEGF siRNA (IL-1β+VEGF siRNA) transfected IVD cells for 24 hours. After stimulation, HMEC-1 cells “ECs” that were plated for 24 hours on 1 μm pore co-culture inserts (ThinCert cell culture insert; Greiner, Monroe, NC) at a density of $3x10^5$ cells were inserted into the wells containing Scram siRNA or VEGF siRNA transfected IVD cells with MCDB media plus 1% FBS. ECs co-cultured with stimulated Scram siRNA treated IVD cells are denoted as $EC_{IL\text{-}1\beta+Scram\ siRNA}$ and ECs co-cultured with stimulated VEGF siRNA treated IVD cells are denoted as $EC_{IL\text{-}1\beta+VEGF\ siRNA}$. Naïve ECs that did not undergo co-culturing were used as a control group. ECs were analyzed via a Matrigel tube formation assay, multiplex (Human

Angiogenesis & Growth Factor 17-Plex, Human Cytokine Panel A 48-Plex, Human MMP/TIMP 13-Plex; MilliporeSigma) and RNA sequencing (Figure S4).

*Co-culture Between IVD cells and SH-SY5Y Cells (NCs)*

SH-SY5Y neuronal cells "NCs" were plated at a density of 1.5 x $10^5$ cells per well in a 6 well plate and cultured for 48 hours prior to the co-culture experiment. For co-culturing, non-stimulated IVD cells "naïve", IL-1β+Scram siRNA, and IL-1β+VEGF siRNA transfected IVD cells were plated on the co-culture insert at a density of 3.0 x $10^5$ cells and co-cultured for 48 hours with NCs. NCs were analyzed via a neurite outgrowth assay and RNA sequencing (Figure S5).

*RNA Preparation and Sequencing*

Post co-culture experiments, the HMEC-1 and SH-SY5Y cells were lysed (TRIzol reagent, Fisher Scientific), and total RNA was extracted (Direct-zol RNA MicroPrep Kit, Zymo Research). Samples were prepared according to library kit manufacturer's protocol, indexed, pooled, and sequenced on an Illumina NovaSeq X Plus. Basecalls and demultiplexing were performed with Illumina's DRAGEN and BCLconvert version 4.2.4 software. RNA-seq reads were then aligned to the Ensembl release 101 primary assembly with STAR version 2.7.9a1. Gene counts were derived from the number of uniquely aligned unambiguous reads by Subread: featureCounts version 2.0.32. Isoform expression of known Ensembl transcripts were quantified with Salmon version 1.5.23. Sequencing performance was assessed for the total number of aligned reads, total number of uniquely aligned reads, and features detected. The ribosomal fraction, known junction saturation, and read distribution over known gene models were quantified with RSeQC version 4.04. All gene counts were then imported into the R/Bioconductor package EdgeR5 and TMM normalization size factors were calculated to adjust for samples for differences in library size. Ribosomal genes and genes not expressed in the smallest group size (minus one sample greater than one count-per-million) were excluded from further analysis. The TMM size factors and the matrix of counts were then imported into the R/Bioconductor package Limma6. Weighted likelihoods based on the observed mean-variance relationship of every gene and sample were then calculated for all samples and the count matrix was transformed to moderated log 2 counts-per-million with Limma's voomWithQualityWeights7. The performance of all genes was assessed with plots of the residual standard deviation of every gene to their average log-count with a robustly fitted trend line of the residuals. Differential expression analysis was then performed to analyze differences between conditions and the results were filtered for only those genes with Benjamini-Hochberg false-discovery rate-adjusted P values less than or equal to 0.05. The raw data files are available on the Gene Expression Omnibus database: GSE294577

*Multiplex Protein Assay*

Conditioned media from naïve ECs and those co-cultured with IL-1β+Scram siRNA and IL-1β+VEGF siRNA transfected IVD cells was collected and analyzed by multiplex protein assay (Human Angiogenesis & Growth Factor 17-Plex, Human Cytokine Panel A 48-Plex, Human MMPTIMP 13-Plex, Eve Technologies, Calgary, Alberta, Canada). Out of range values or results < 2 pg/ml were not included.

*Matrigel Tube Formation Assay*

To determine the vessel formation capacity between ECs cultured with conditioned media (CM) collected from IL-1β+Scram siRNA versus IL-1β+VEGF siRNA transfected IVD cells, a Matrigel tube formation assay was performed. Matrigel matrix (356231; Corning Inc., Corning, NY), a gelatinous protein mixture that mimics the extracellular matrix, was used to coat a 24-well plate following the manufacturer's protocol. HMEC-1 cells at the density of 3 x $10^4$ cells per well were suspended in the CM collected from IL-1β+Scram siRNA or IL-1β+VEGF siRNA transfected IVD cells for 48 hours and plated onto each Matrigel coated well. Images were acquired at 6 hours and 12 hours post plating via a Nikon Eclipse Ti2 inverted scope at 10×. To assess the ability of tube formation by each conditioned media and time point, the numbers of closed loops were counted[22].

*Neurite Outgrowth Assay*

The number and length of neurites from NCs co-cultured with non-stimulated IVD cells "naïve", IL-1β+Scram siRNA, or IL-1β+VEGF siRNA transfected IVD cells were compared. Neurite outgrowth was quantified using ImageJ software. For each neuron, neurite number and length were counted manually. Neurite length was measured from the edge of the cell body to the tip of the neurite. The average number of neurites, and average length and total neurite lengths, in microns, were calculated for each treatment group.

Supplemental Tables and Figures

| Assay | Sex M/F | Tamoxifen +/- | Naïve/Sham |
|---|---|---|---|
| Von Frey | 0.94 | 0.63 | 0.66 |
| Grip Strength | 0.09 | 0.92 | 0.07 |
| Inverted Wire Hang | 0.65 | 0.33 | 0.64 |
| Cold Sensitivity | 0.52 | 0.09 | 0.40 |
| Heat Sensitivity | 0.44 | 0.47 | 0.57 |
| Rotarod | 0.34 | 0.43 | 0.25 |
| Distance (Open Field) | 0.33 | 0.27 | 0.18 |
| Rearing (Open Field) | 0.25 | 0.32 | 0.59 |
| Neurite Length | 0.08 | [redacted] | [redacted] |
| Vessel Length | 0.19 | [redacted] | [redacted] |
| Neurite Depth | 0.51 | [redacted] | [redacted] |
| Vessel Depth | 0.15 | [redacted] | [redacted] |

**Table S1: Behavioral and Neurovascular Comparisons for Sex, Tamoxifen, and Naïve/Sham.** Assays with no evidence of differences between genotypes at any postoperative time point following IVD injury. The respective p-values are shown for comparisons between Male and Female, Tamoxifen +/-, and Naïve and Sham surgery for the behavioral, neuronal, and vascular measurements. P-values from Linear Mixed Effects model.

| Group | Factor | Scrambled siRNA | VEGFA siRNA | p-value (paired) |
|---|---|---|---|---|
| Angiogenesis | Angiopoietin-2 | 29.1±2.1 (pg/ml) | 28.9±2.4 (pg/ml) | 0.53 |
| Angiogenesis | Endoglin | 84.8±10 | 92.9±5.9 | 0.06 |
| Angiogenesis | Endothelin-1 | 337.3±42.5 | 336.7±34.9 | 0.84 |
| Angiogenesis | Follistatin | 436.8±485.1 | 915.9±1105.5 | 0.12 |
| Angiogenesis | Leptin | 70.4±11 | 70.4±8.4 | 0.79 |
| Angiogenesis | PlGF | 127.9±4 | 126.2±7.6 | 0.88 |
| MMP | MMP2 | 8775±1081.8 | 9309.3±1816.7 | 0.55 |
| MMP | MMP7 | 613.5±45.3 | 759.9±120.3 | 0.24 |
| MMP | MMP8 | 138.6±116.4 | 239.4±120 | 0.30 |
| MMP | MMP9 | 163±30.5 | 249.7±44.1 | 0.05 |
| MMP | MMP12 | 582.4±65.1 | 604.9±59.9 | 0.60 |
| MMP | TIMP1 | 100638.6±13530.7 | 89032.7±6718.9 | 0.13 |
| MMP | TIMP2 | 30098.3±2192.1 | 31829.1±3278.2 | 0.07 |
| MMP | TIMP3 | 3770.7±1298.2 | 2823.7±677.1 | 0.42 |
| MMP | TIMP4 | 56.8±6.4 | 54.1±3.1 | 0.55 |
| Cytokines | EGF | 3.2±0.8 | 3.5±0.7 | 0.64 |
| Cytokines | CCL11 | 7.7±0.6 | 7.6±0.7 | 0.89 |
| Cytokines | FGF2 | 187±72.4 | 200.5±58.4 | 0.23 |
| Cytokines | FLT3L | 0.6±0.1 | 0.6±0.1 | 0.74 |
| Cytokines | M-CSF | 246.1±74.1 | 222.9±61 | 0.60 |
| Cytokines | IL-6 | 2481.3±183 | 2518±201.4 | 0.94 |
| Cytokines | IL-8 | 14958±894.4 | 16325.9±275.8 | 0.09 |
| Cytokines | CCL2 | 3441.9±1491.7 | 3898.9±1275.8 | 0.18 |
| Cytokines | PDGF-BB | 32.9±7.5 | 20.9±5.8 | 0.45 |

**Table S2. Panel of measured cytokines produced by the EC after co-culturing with the VEGFA-silenced IL-1β–stimulated human IVD cells.**

| | | Pfirrmann Grade | | | | | | | | | | |
|---|---|---|---|---|---|---|---|---|---|---|---|---|
| **Age** | **Sex** | **T12/L1** | **L1/2** | **L2/3** | **L3/4** | **L4/5** | **L5/S1** | **Mean** | **STD** | **Endothelial Cell (RNA/ Multiplex/ ELISA)** | **Matrigel Outgrowth Assays** | **Neural Cell (RNA/Neurite)** |
| 70 | F | | 3 | 3 | 4 | 4 | 4 | 3.6 | 0.5 | | X | |
| 73 | F | | | 3 | 3 | 4 | 5 | 3.8 | 1.0 | X | | |
| 73 | M | | | 3 | 3 | 3 | 4 | 3.3 | 0.5 | | | X |
| 66 | F | 3 | 3 | 3 | 5 | 5 | | 3.8 | 1.1 | X | X | |
| 50 | M | | | | | 3 | 4 | 3.5 | 0.7 | X | | |
| 76 | F | | 2 | 3 | 3 | 4 | | 3.0 | 0.8 | | | X |
| 68 | F | | | | | 3 | 3 | 3.0 | 0.0 | | | X |
| 72 | M | 3 | 3 | 4 | 4 | 4 | 5 | 3.8 | 0.8 | | X | |

**Table S3: Patients' Demographics for Intervertebral Disc Cells:** IVD cells were isolated from eight patients removed during elective surgical procedures for degenerative spinal disease. Pfirrmann Grades for T12/S1 are shown along with experimental distributions for each patient sample.

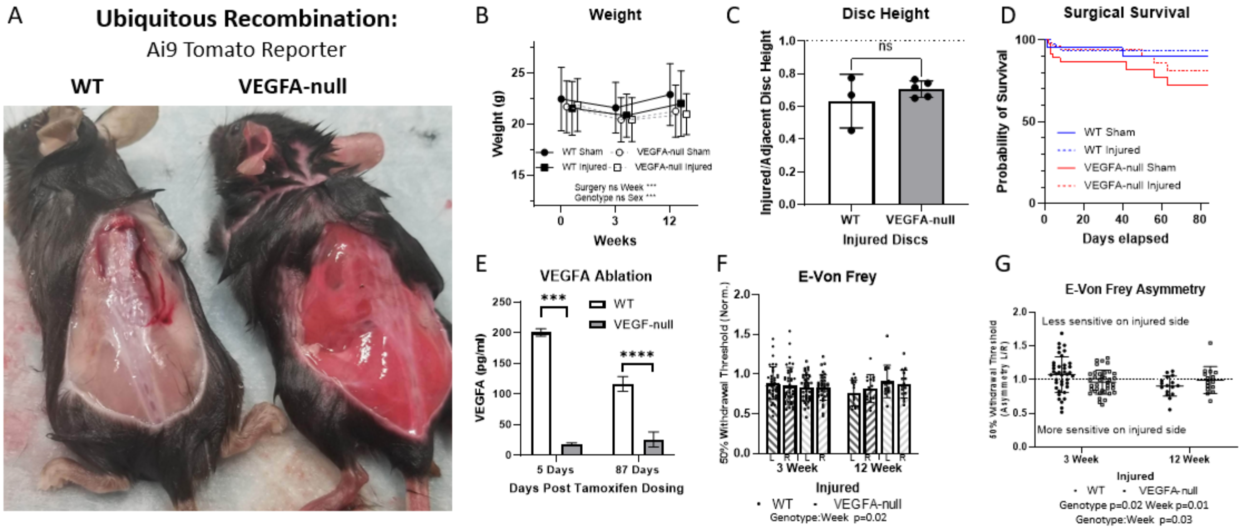


**Figure S1 VEGFA ablation is highly efficient and does not pose health issues in the post-natal animal.** A) Ubiquitous recombination in a WT and VEGFA-null showing the pink WT and red VEGFA-null Ai9 tdTomato reporter B) Mouse body weights at 0 (Baseline), 3 and 12 weeks remain similar between groups irrespective of genotype or surgery type C) Disc height ratio of injured/adjacent control disc height showing a similar decrease in heights following injury D) Surgical survival curves show no significant differences between genotype or surgery E) VEGFA ablation confirmation after initial ablation and 12 weeks from FSU media after TNFα stimulation ELISA showing a sustained ablation over time F) E-Von Frey shows that WT injured sensitivity increased over time (decreased withdrawal threshold) while VEGFA-null decreased over time (increased withdrawal threshold) G) E-Von Frey asymmetry signifies an imbalance in WT mice that fluctuates over time while VEGFA-null mice remained nearer to symmetry (dotted line). Data are presented as mean ± standard deviation.

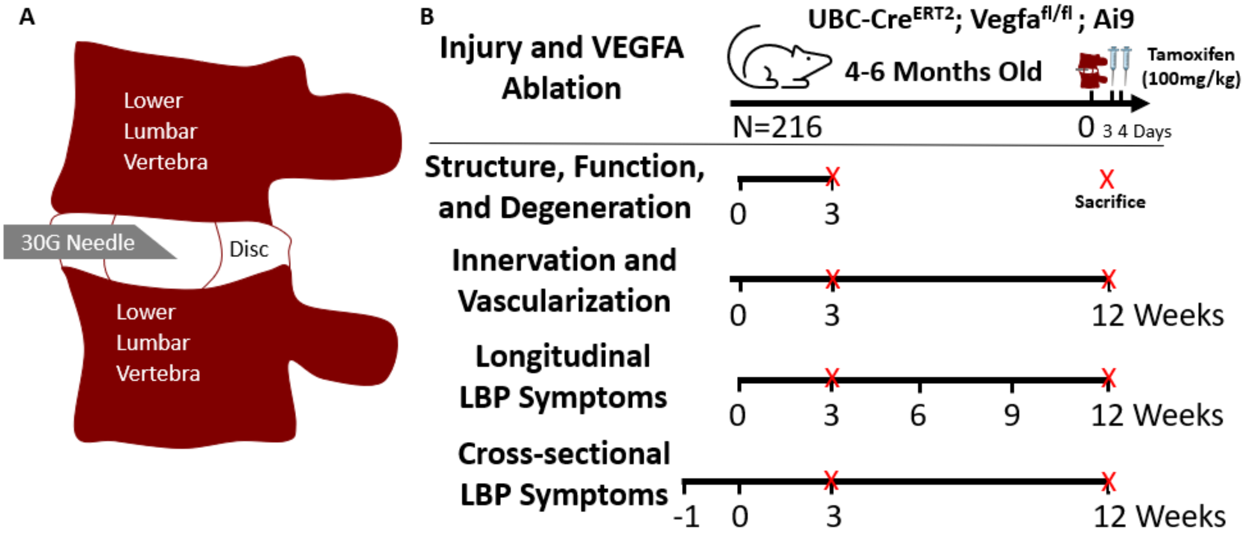


**Figure S2: Experimental Design** A) Lower lumbar disc puncture model showing the relative scale of a 30G needle and the depth of injury B) Experimental design for various outcomes including injury at time=0 with VEGFA ablation at 3-4 days post-op, and acute and chronic terminal time points at 3 and 12 weeks. Baseline behavior within one week or surgery normalized behavioral outcomes. N=216 total mice.

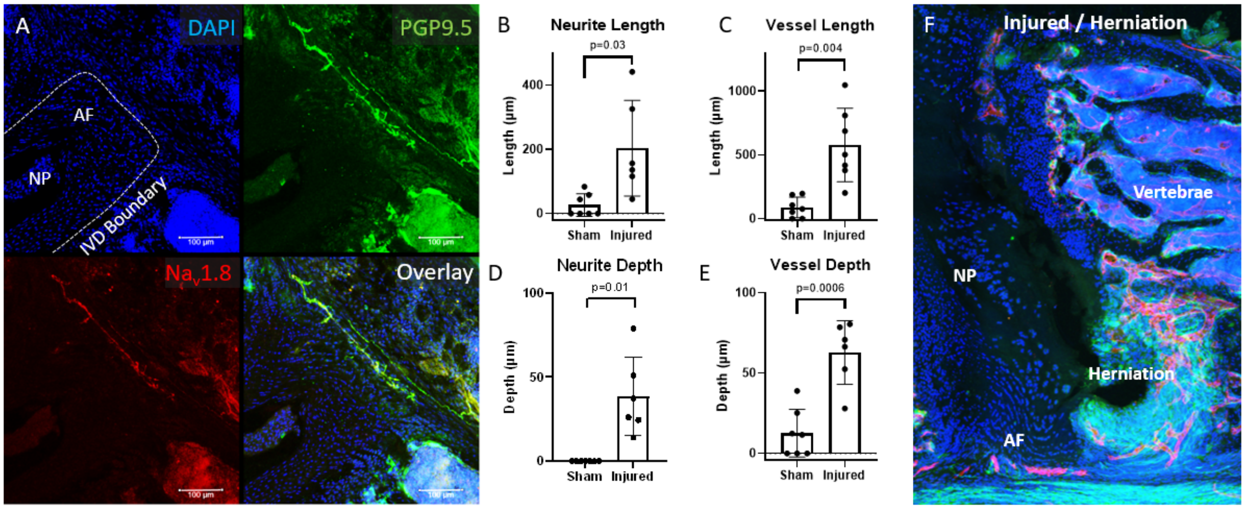


**Figure S3: De novo PGP9.5+ structures in the IVD following injury are sensory neurons.** A) Validation of PGP9.5 IHC with a $Na_v1.8$- tdTomato reporter showing the coincidence of the more specific sensory $Na_v1.8$ with pan neuronal PGP9.5. $Na_v1.8$+ features accounted for 94% of the $PGP9.5^+$ features with a spatial colocalization correlation of $R = 0.93$. B-E) Acute (3 week) neurite length and depth and vessel length and depth data showing the increased lengths associated with injured discs F) IHC of a herniated disc at 3 weeks showing diffuse nonspecific green PGP9.5 staining and red Endomucin vessels. Data are presented as mean ± standard deviation.

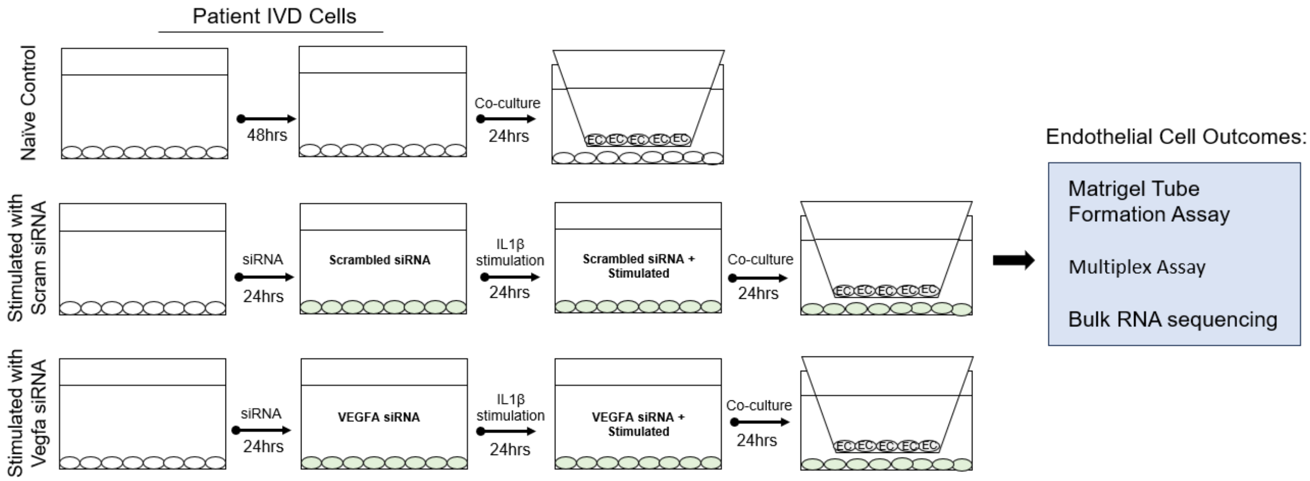


**Figure S4. Schematic drawing of Co-culture between IVD cells and HMEC-1(ECs)** IVD cells were plated on a 6 well plate left as naïve or transfected with Scrambled control siRNA (Scram siRNA) or VEGFA specific siRNA (VEGFA siRNA) for 24 hours. Cells were then treated with 1 ng/mL of IL-1β to stimulate the Scram siRNA and VEGF siRNA transfected IVD cells (IL-1β+Scram siRNA or siRNA IL-1β+VEGF siRNA, respectively) for 24 hours but the naïve IVD cells were left unstimulated. After stimulation, 1 μm pore co-culture inserts plated with ECs were inserted into each well for direct coculturing for 24 hours.

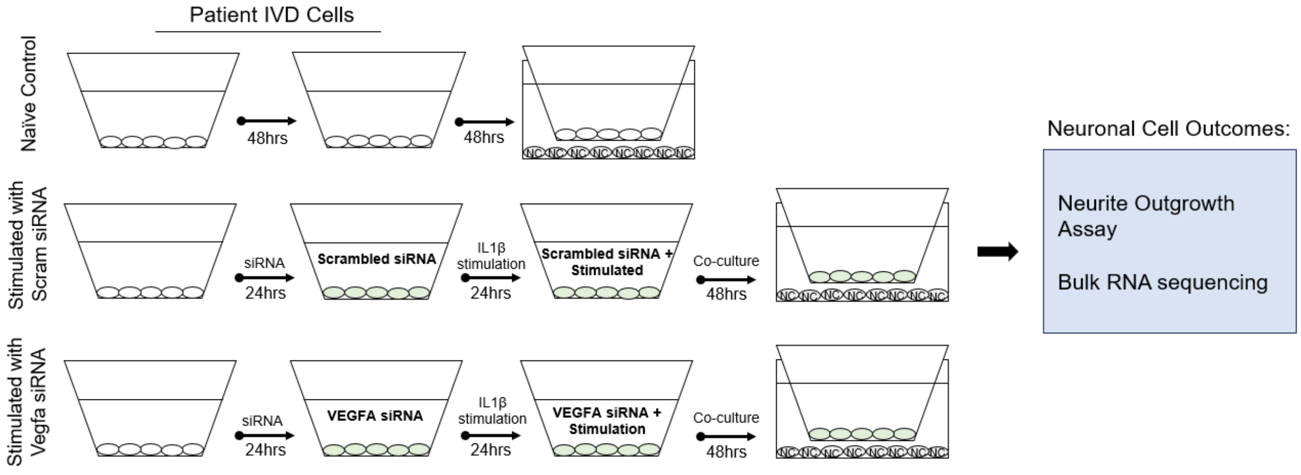


**Figure S5. Schematic drawing of Co-culture between IVD cells and SH-SY5Y cells (NCs).** NCs were plated on a 6 well plate and cocultured with 1 µm pore co-culture inserts plated with IVD cells 48 hours. IVD cells were either left as naïve or transfected with Scrambled control siRNA (Scram siRNA) or VEGFA specific siRNA (VEGFA siRNA) for 24 hours. The naïve IVD cells were left unstimulated and the Scram siRNA and VEGF siRNA transfected IVD cells were stimulated with 1 ng/mL of IL-1β (IL-1β+Scram siRNA or siRNA IL-1β+VEGF siRNA, respectively) for 24 hours prior to co-culture.

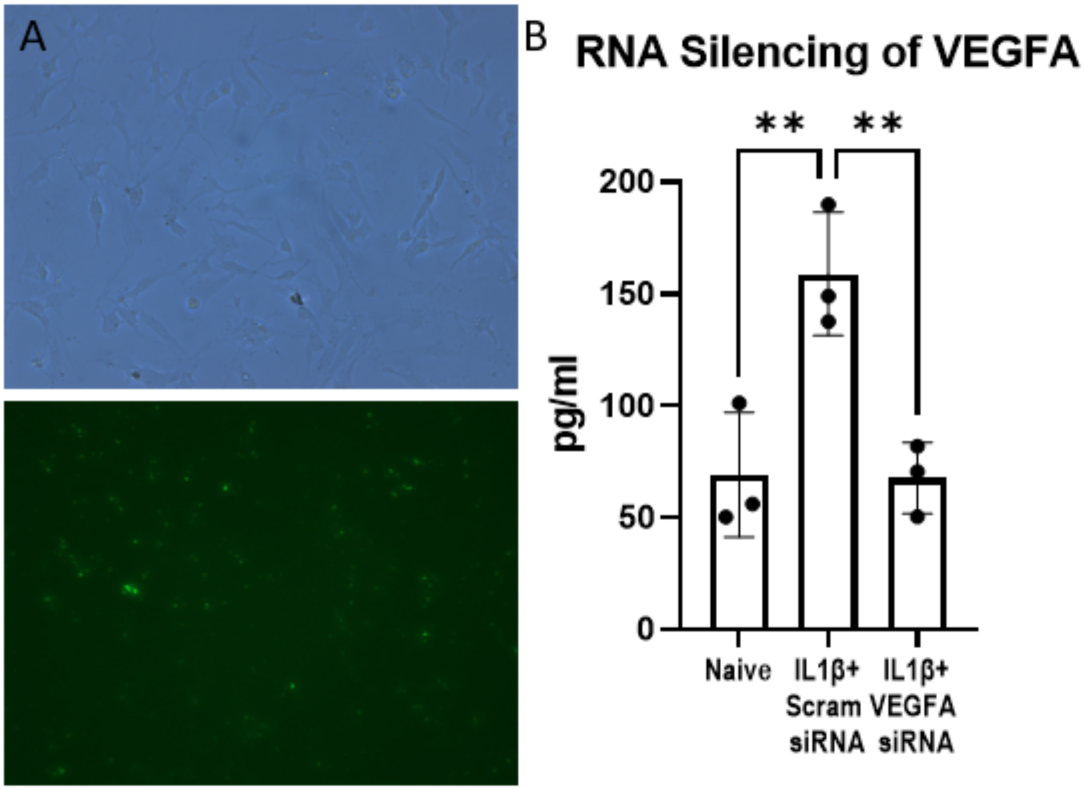


**Figure S6. Validation of VEGFA-siRNA Transfection in IVD Cells** A) Control siRNA (Fluorescein Conjugate)-A revealed efficient transfection of cationic lipid-based transfection reagents in IVD cells. B) Stimulated IVD cells by IL-1β with Scrambled siRNA show an increase in the VEGFA secretion compared to naïve IVD cells ($p=0.04$). Stimulated IVD cells with IL-1β after being transfected with VEGFA siRNA showed a significant silencing (57.4%) of VEGFA secretion compared to Scram siRNA ($p=0.03$).
